# A structure-informed deep learning framework for modeling TCR-peptide-HLA interactions

**DOI:** 10.64898/2026.03.31.715361

**Authors:** Kai Cao, Rui Li, Martin Stražar, Eric M. Brown, Phuong N.U. Nguyen, Marie-Madlen Pust, Jihye Park, Daniel B. Graham, Orr Ashenberg, Caroline Uhler, Ramnik J. Xavier

**Affiliations:** Broad Institute of MIT and Harvard, Cambridge, MA, United States; Department of Molecular Biology, Massachusetts General Hospital and Harvard Medical School, Boston, MA, United States; Klarman Cell Observatory, Broad Institute of MIT and Harvard, Cambridge, MA, United States; Massachusetts Institute of Technology, Cambridge, MA, United States; Center for the Study of Inflammatory Bowel Disease, Massachusetts General Hospital, Boston, MA, United States

## Abstract

The interaction between T cell receptors (TCRs), peptides, and human leukocyte antigens (HLAs) underlies antigen-specific T cell immunity. Despite substantial advances in peptide– HLA presentation prediction, accurate modeling of coupled TCR–peptide–HLA recognition remains underdeveloped, limiting applications such as TCR and neoepitope prioritization in cancer and antigen identification in autoimmunity. Here we present StriMap, a unified framework for predicting TCR–peptide–HLA interactions by integrating physicochemical, sequence-context, and structural features at recognition interfaces. StriMap achieves state-of-the-art performance with improved generalizability and enables applications in both cancer and autoimmunity. As a case study in ankylosing spondylitis (AS), we screened 13 million peptides derived from 43,241 bacterial proteins and identified candidate molecular mimics that were experimentally validated to activate T cells expressing an AS-associated TCR. Notably, a top validated peptide was enriched in patients with inflammatory bowel disease (IBD), suggesting potential shared microbial triggers between AS and IBD. Overall, StriMap provides a generalizable framework for rational immunotherapy design and for dissecting antigenic drivers of autoimmunity.

## Introduction

T cell–mediated immunity relies on the recognition of antigenic peptides presented by human leukocyte antigen (HLA) molecules through T cell receptors (TCRs) expressed on the surface of T cells^1^. This HLA-restricted recognition of peptide epitopes is central to adaptive immunity, enabling the detection of infected or transformed cells while maintaining tolerance to self. This same recognition system, which enables broad immune surveillance, can also determine disease trajectories when dysregulated or redirected. Cancer and autoimmunity represent two challenging and important problems in which antigen-specific T cell recognition is central to both pathogenesis and therapy. In cancer, the ability to prioritize TCRs and neoepitopes underpins personalized immunotherapies and vaccines^2,3^. In autoimmunity, identifying disease-associated antigens and potential molecular mimics can reveal causal triggers and enable antigen-specific interventions^4,5^.

While several experimental platforms can detect interactions between TCRs and peptide–HLA (pHLA) complexes^6–8^, they remain time-consuming and expensive, limiting coverage of TCR or peptide diversity. These limitations motivate the need for computational approaches^9^. Despite major advances in computational prediction of pHLA presentation^10–13^, systematically mapping downstream TCR–pHLA recognition remains underdeveloped^9^. This difficulty reflects the combinatorial complexity of the interaction space, the structural plasticity of recognition, and the limited and biased nature of available experimental data, as reflected in the most recent benchmark studies^14–16^. Deep learning has begun to be used to model these processes, yet often relies on sequence-based representations and treats pHLA presentation and TCR–pHLA recognition as independent tasks^17–24^, thereby limiting generalizability beyond the training distribution by failing to capture their conditional dependence and shared structural constraints. Recent efforts have begun to incorporate structural information and multimodal representations, such as structure-based graph neural network approaches (e.g., STAG^25^) and hybrid sequence– structure models integrating protein language models (e.g., STAG-LLM^26^), demonstrating the value of structural features in modeling TCR–pHLA interactions. However, a unified framework that explicitly links peptide–HLA presentation with downstream TCR recognition remains lacking. Bridging this gap requires a framework that sequentially models peptide–HLA presentation and TCR recognition to better reflect the underlying biological process, while integrating complementary sequence, structural, and contextual features.

To address this gap, we developed StriMap (<u>S</u>tructure-informed TRi-molecular Interaction Mapping), a unified deep learning framework that models antigen presentation and TCR recognition as a coupled biological process, integrating complementary sequence, structural, and contextual features to capture the determinants of immune recognition. Across diverse public datasets curated from multiple independent studies^11,12,14–16,21,22,24^, StriMap consistently achieves state-of-the-art performance when benchmarked against leading HLA presentation and TCR-specificity models. We further demonstrate the utility of StriMap in both cancer and autoimmunity. In cancer, StriMap enables TCR-centric prioritization of recurrent neoepitopes and supports neoepitope selection for personalized vaccine design. In autoimmunity, we applied StriMap to investigate the link between HLA-B27–bound bacterial peptides and TCRs containing the conserved TRBV9 motif associated with ankylosing spondylitis (AS). Leveraging a small amount of recently published disease-specific data together with large-scale public TCR– pHLA datasets, we trained the model to screen 13 million peptides derived from 43,241 bacterial proteins in gut bacterial strains linked to AS, ranking candidates by predicted binding scores. Notably, top-ranked peptides were experimentally validated to activate T cells expressing a representative TRBV9 TCR. Matched metagenomic and metatranscriptomic analyses revealed that one validated peptide from *Streptococcus* strains was enriched in patients with inflammatory bowel disease (IBD), suggesting a shared antigenic trigger between the two autoimmune diseases. Together, these results highlight StriMap as a generalizable framework for linking antigen presentation to TCR recognition across cancer and immune-mediated disease. To facilitate broad use, we provide an accessible web portal (www.strimap.com) to run predictions and train models on user-provided data.

## Results

### StriMap overview

To systematically model the tri-molecular interaction underlying T cell-mediated immunity (Fig. 1a), we developed StriMap, a unified deep learning framework that integrates physicochemical, sequence-context, and structural features. At the core of StriMap is the Sequence and Structure Feature Extractor (SSFE), a multimodal encoder that learns biophysical determinants of molecular recognition (Fig. 1b). For each input (peptide, HLA, and paired TCR α/β chains), the SSFE generates a comprehensive representation by fusing three complementary channels: (*i*) residue-level physicochemical features^27^ (Table S1); (*ii*) evolutionary sequence context from pretrained protein language models (e.g., ESM2^28^, ProtT5^29^); and (*iii*) 3D structural features predicted by ESMFold^28^ and processed via an equivariant graph neural network (EGNN)^30^ to encode spatial geometries (see Methods). We further introduce a bilinear attention mechanism^31^ to model residue–residue interactions between molecular components (Fig. 1c). To reflect the hierarchical nature of immune recognition, StriMap employs a coupled architecture in which peptide–HLA presentation is first modeled and subsequently used to condition TCR recognition on the underlying peptide–HLA landscape (Fig. 1d–e), thereby enforcing biological consistency. To address the scarcity of validated negative TCR–pHLA pairs, we introduce dynamic and mutation-aware negative sampling to improve discrimination between true binders and near-neighbor decoys. Together, these architectural and training innovations enable StriMap to function as a unified framework for TCR or neoepitope prioritization in cancer immunotherapy and the identification of disease-associated antigens in autoimmunity (Fig. 1f), which we systematically evaluate in the following sections.

**Fig. 1.**
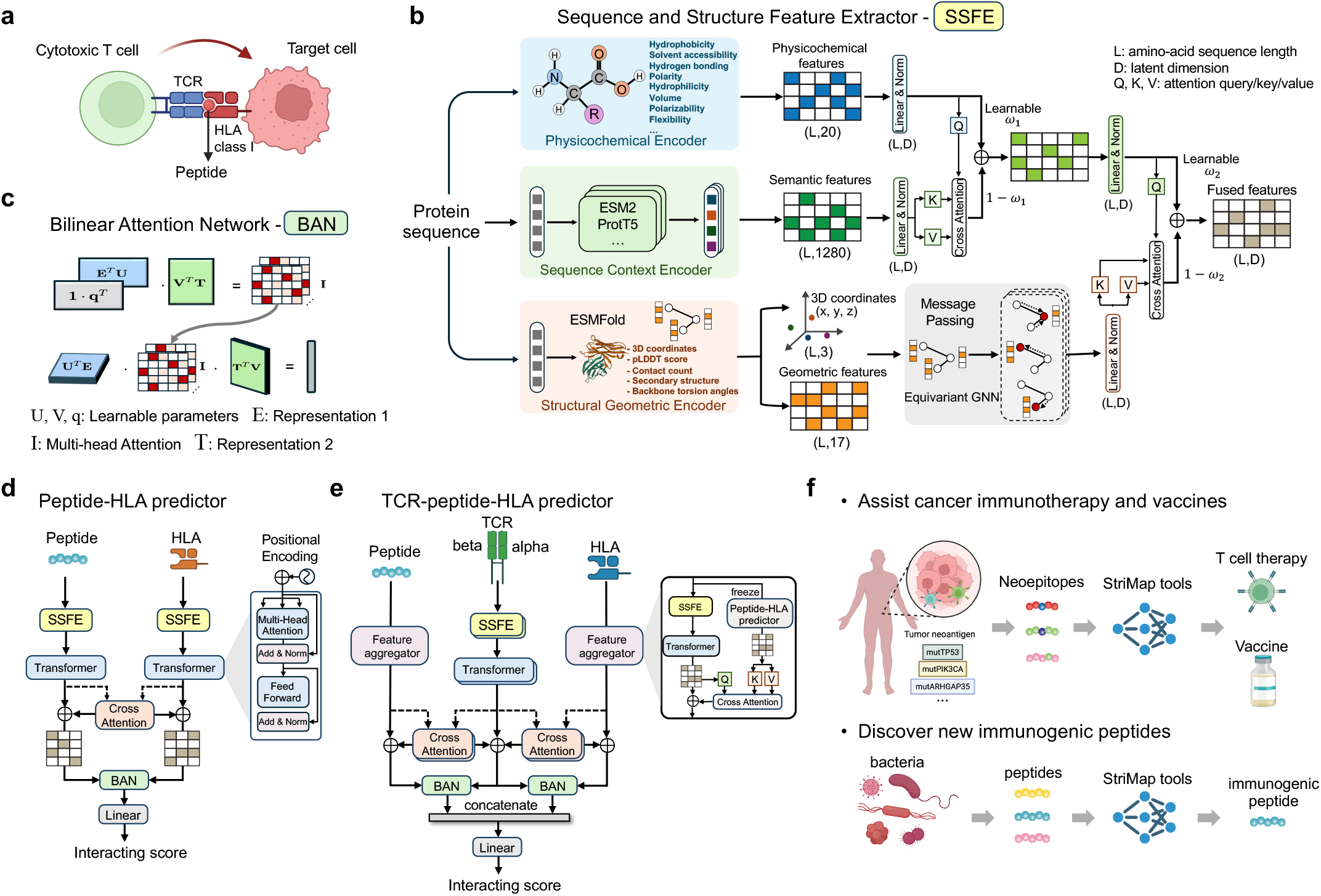
Overview of StriMap framework. **(a)** Biological context of T cell-mediated immune recognition. Cytotoxic T cells recognize target cells through the interaction between the T cell receptor (TCR) and peptide-HLA (pHLA) complexes presented on the surface of target cells. This tri-molecular interaction underlies antigen-specific immune surveillance and forms the biological basis of StriMap.**(b)** The architecture of the Sequence and Structure Feature Extractor (SSFE). Given a protein amino-acid sequence, SSFE integrates three complementary encoders: i) a physicochemical encoder that computes residue-level physicochemical features; ii) a sequence-based context encoder (e.g., ESM2 or ProtT5) that provides contextual sequence embeddings; and iii) a structural-derived geometric encoder (e.g., ESMFold) that generates 3D structural representations. Structural information is further processed through a multi-layer equivariant graph neural network (EGNN). All features are fused using gated cross-attention and layer-normalization to produce a unified representation.**(c)** The bilinear attention network (BAN), which models pairwise interactions between two representations using learnable bilinear projections to generate multi-head attention maps.**(d)** Architecture of the peptide-HLA interaction predictor. Peptide and HLA sequences are independently encoded by SSFE and processed through transformer blocks, followed by gated cross-attention and BAN fusion. The fused representation is fed into a linear layer to produce an interaction score between 0-1.**(e)** Architecture of the TCR-peptide-HLA interaction predictor. TCR α/β chains, peptide, and HLA sequences are independently encoded by SSFE and then passed through a feature aggregator module that incorporates and leverages the embedding representations learned from the peptide-HLA predictor. The aggregated features are further integrated via gated cross-attention and BAN, and the fused representation is mapped through a linear layer to produce an interaction score between 0-1.**(f)** Downstream applications of StriMap. StriMap enables systematic modeling of peptide-HLA and TCR-peptide-HLA interactions, facilitating the prioritization of immunogenic neoepitopes for cancer immunotherapy and vaccine development, as well as the discovery of pathogenic or immunodominant epitopes.

### StriMap achieves improved performance across pHLA and TCR–pHLA benchmarks

We evaluated StriMap against recently published state-of-the-art models on two related tasks— pHLA presentation prediction and TCR–pHLA interaction prediction—using benchmark datasets curated from independent prior studies with published train/test splits to enable reproducible comparison (Table S2–S3). Baselines were taken from the original studies to avoid implementation biases. We considered in-distribution (ID) testing, where test examples are drawn from the same underlying distribution as training; distribution-shifted (DS) testing across different studies or cohorts (dataset-level shift); and epitope-level out-of-distribution (OOD) generalization to epitopes not seen in training, forming an increasingly stringent hierarchy from ID to DS to OOD.

Across multiple pHLA presentation benchmarks^11,12,21^, StriMap achieved top overall performance and remained robust under increasingly stringent ID/DS/OOD settings (Fig. 2a), with extended metric-level benchmarking results summarized in Fig. S1. Performance gains were also consistent across individual HLA alleles: on the Que et al. dataset (ID)^21^, introduced alongside deepAntigen, StriMap improved per-allele AUPRC relative to deepAntigen^21^ (Fig. 2b), and broader per-allele comparisons against additional state-of-the-art predictors further supported consistent improvements across HLA classes (Fig. S2). StriMap also consistently surpassed NetMHCpan^10^ across all evaluations, underscoring robust gains over a widely used reference method in the literature. These gains translated to tumor neoantigen prioritization: under the same evaluation protocol used in deepAntigen, StriMap achieved higher true discovery rates across ranking thresholds on the TESLA (Tumor Neoantigen Selection Alliance)^32^ benchmark (Fig. 2c), and similarly improved performance on the independent CEDAR (Cancer Epitope Database and Analysis Resource)^33^ benchmark compared with NetMHCpan^10^ (Fig. S3). Notably, while performance differences are modest at lower ranking thresholds, StriMap shows consistently stronger enrichment among top-ranked candidates than NetMHCpan, which is most relevant for practical neoantigen prioritization where only a limited number of candidates can be experimentally tested.

**Fig. 2.**
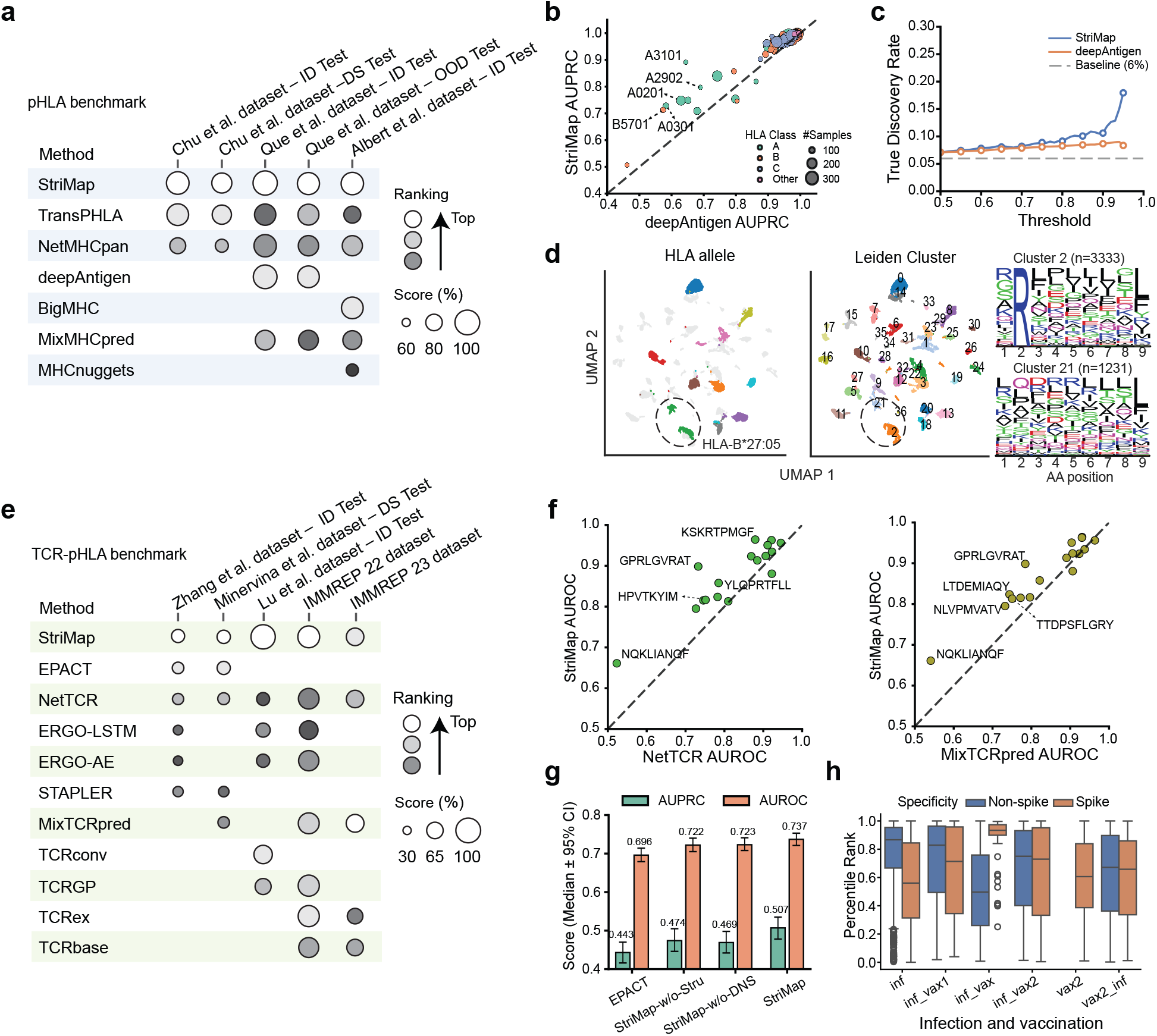
StriMap achieves robust prediction of pHLA presentation and TCR-pHLA recognition. **(a)** Summary of pHLA presentation prediction performance across multiple benchmark datasets curated from independent prior studies. Methods are evaluated under in-distribution (ID), distribution-shifted (DS), and out-of-distribution (OOD) settings as defined by the original publications. Circle size indicates relative performance, and color denotes the corresponding dataset. Rankings are determined based on overall performance, with lighter color intensity indicating higher rankings within the dataset; white denotes the top-ranked method.**(b)** Comparison of StriMap and deepAntigen performance (AUPRC) across individual HLA alleles on the Que et al. dataset (ID). Each point represents one HLA allele, with color indicating HLA class and size denoting the number of peptide samples. The dashed diagonal line indicates equal performance.**(c)** True discovery rate (TDR) as a function of ranking threshold on the TESLA neoantigen benchmark. StriMap is compared with deepAntigen under the same evaluation protocol. The dashed line indicates the baseline discovery rate.**(d)** UMAP visualization of peptide embeddings, colored by 10 most abundant HLA alleles (left) and Leiden clusters (right). Two distinct clusters (dashed ellipse) corresponding to HLA-B*27:05 exhibit divergent peptide-binding motifs, as illustrated by sequence logos.**(e)** Summary of TCR–pHLA interaction prediction performance across multiple benchmark datasets. Methods are evaluated under ID and DS settings as defined by the original publications. Circle size indicates normalized performance score, and color denotes the dataset. Rankings are determined based on overall performance, with color intensity encoding ranking; white denotes the top-ranked method.**(f)** Per-peptide AUROC comparison between StriMap and NetTCR and MixTCRpred on the Zhang et al. dataset DS benchmark. Each point corresponds to one peptide epitope. The dashed diagonal line indicates equal performance.**(g)** Comparison of AUROC and AUPRC performance for StriMap and baseline methods under the same evaluation protocol, including ablation variants. Bars indicate median performance across test sets, with error bars denoting 95% confidence intervals. StriMap-w/o-Stru denotes StriMap without structural information, and StriMap-w/o-DNS denotes StriMap without dynamic negative sampling.**(h)** Percentile binding rank distributions of spike-specific and non-spike-specific TCRs across six cohorts with distinct SARS-CoV-2 infection and vaccination histories. Cohorts include recovered individuals sampled post-infection (inf), recovered individuals after one vaccine dose (inf_vax1), immediately after two doses (inf_vax), post-vaccination recovery (inf_vax2), vaccinated naïve individuals (vax2), and breakthrough infection cases (vax2_inf).

To probe what StriMap learns beyond aggregate accuracy, we trained a pHLA model on a comprehensive dataset assembled from multiple sources (Table S2). This dataset was allele-imbalanced (Fig. S4). Visualization of the learned embedding space revealed a biologically meaningful organization, where the cluster separation reflected not only the individual HLA alleles but also the fine-grained heterogeneity within each allele, determined by distinct peptide-binding properties (Fig. 2d; Fig. S5). For example, in HLA-B*27:05, peptides segregated into two major Leiden clusters (2 and 21) with distinct motifs: cluster 2 recapitulated the canonical preference for Arg at position 2 and Leu at the C terminus, whereas cluster 21 showed non-canonical preferences at position 2, indicating sub-allelic structure beyond a single consensus motif. Complementing embedding-level structure, attention-based analyses further revealed allele-specific binding-position profiles and distinct interaction patterns between peptide residues and HLA contact residues, including differences between closely related alleles such as HLA-A*02:01 and HLA-A*02:03 (Fig. S6). Finally, systematic saturation mutagenesis on a well-characterized influenza epitope (GILGFVFTL) demonstrated allele- and position-specific sensitivity in predicted binding scores (Fig. S7), supporting that StriMap captures biologically meaningful constraints underlying peptide–HLA binding.

We next evaluated StriMap on the more important task of TCR–pHLA interaction prediction across multiple independent benchmarks^6,14–16,34^. For training, negative examples were generated by random TCR and pHLA shuffling, consistent with prior benchmarking practices^35^. We additionally incorporated dynamic negative sampling (DNS), regenerating negatives each epoch of model training to expand the effective negative space. Overall, StriMap achieved strong performance under both ID and DS settings (Fig. 2e), with per-peptide comparison on Zhang et al. (ID), Minervina et. al (DS) and IMMREP22 (Fig. S8–S10); Fig. 2f highlights the IMMREP22 per-peptide AUROC comparison, where StriMap outperformed NetTCR^17^ and MixTCRpred^23^, two widely used reference methods. Ablation analyses further showed that removing the structural module and/or DNS degraded AUROC/AUPRC relative to the full model (Fig. 2g). We note that the Lu et al. benchmark^16^ adopts peptide-permutation negatives that preserve TCR– HLA pairs but scramble peptide identity (Fig. S11). Under this scheme, negatives can be separated largely via peptide–HLA compatibility rather than true TCR-specific recognition, potentially inflating performance. Because StriMap jointly models pHLA presentation and TCR– pHLA recognition, it can exploit peptide–HLA compatibility signals and achieves substantially higher performance under this protocol (Fig. S11). In contrast, methods that ignore HLA information or model TCR–pHLA interactions in isolation are structurally disadvantaged.

To assess generalization across clinical cohorts, we examined the SARS-CoV-2–related TCR– pHLA interactions in the Minervina et al. dataset^34^, comprising 3,570 TCRs across 14 peptides (5 spike and 9 non-spike). Cohorts included recovered individuals’ post-infection (inf), recovered after one vaccine dose (inf_vax1), sampled immediately after two doses (inf_vax), post-vaccination recovery (inf_vax2), vaccinated naïve individuals (vax2), and breakthrough infections (vax2_inf) (Fig. 2h). We predicted the TCR binding score for each spike or non-spike peptide across all categories, and summarized the results as percentile binding rank distributions (Fig. 2h). Vaccination-associated cohorts (“vax”) are expected to mount elevated spike-specific T cell responses following spike immunization; indeed we found more favorable predicted binding ranks for spike-specific TCRs. This effect peaked in the inf_vax cohort, where spike-specific TCRs achieved the highest predicted binding-rank percentiles, consistent with recent vaccination driving expansion and activation of spike-specific clonotypes. In contrast, at later time points (inf_vax2) and in breakthrough infections (vax2_inf), this separation was attenuated, likely reflecting contraction of vaccine-induced responses and diversification of the T cell repertoire following additional antigen exposures.

We further assessed StriMap on the IMMREP23 challenge^15^, where participants could incorporate external training data beyond the official dataset, and the exact data sources used by different methods are not fully disclosed. We compared three training strategies: (*i*) training only on the organizer-provided data, (*ii*) training on a comprehensive dataset assembled from diverse public studies^14,15,22,36–38^ (Fig. S12), and (*iii*) an ensemble combining multiple negative sampling strategies (Fig. S13a). With the ensemble strategy, StriMap ranked third with AUC0.1=0.6943 (Fig. S13b). Notably, the top two methods (IMW-DETECT and MixTCRpred_s2) leveraged undisclosed in-house data, complicating attribution of gains to architecture versus data. We further evaluated StriMap in a zero-shot (OOD) setting with test epitopes absent from training; despite only three unseen epitopes, performance exceeded random baselines, indicating partial generalization, but remained limited, potentially due to substantial epitope divergence (Fig. S13c). These results suggest that few-shot learning is often more reliable in practice, while still motivating downstream use cases such as cancer TCR and neoepitope prioritization.

### TCR and neoepitope prioritization for cancer immunotherapy

Somatic mutations in cancer can generate neoepitopes that are processed and presented by HLA molecules and can be recognized by T cells (Fig. 3a), providing a key rationale for T cell-based immunotherapies. To demonstrate the translational utility, we applied StriMap to two clinically relevant cancer immunotherapy tasks: (*i*) TCR-centric prioritization for recurrent mutant peptide-HLA pairs in the context of adoptive T cell therapy, and (*ii*) neoepitope prioritization for personalized cancer vaccine design (Fig. 3b). These two settings reflect common decision points in T cell–based immunotherapy, where only a small number of candidates can be experimentally evaluated.

**Fig. 3.**
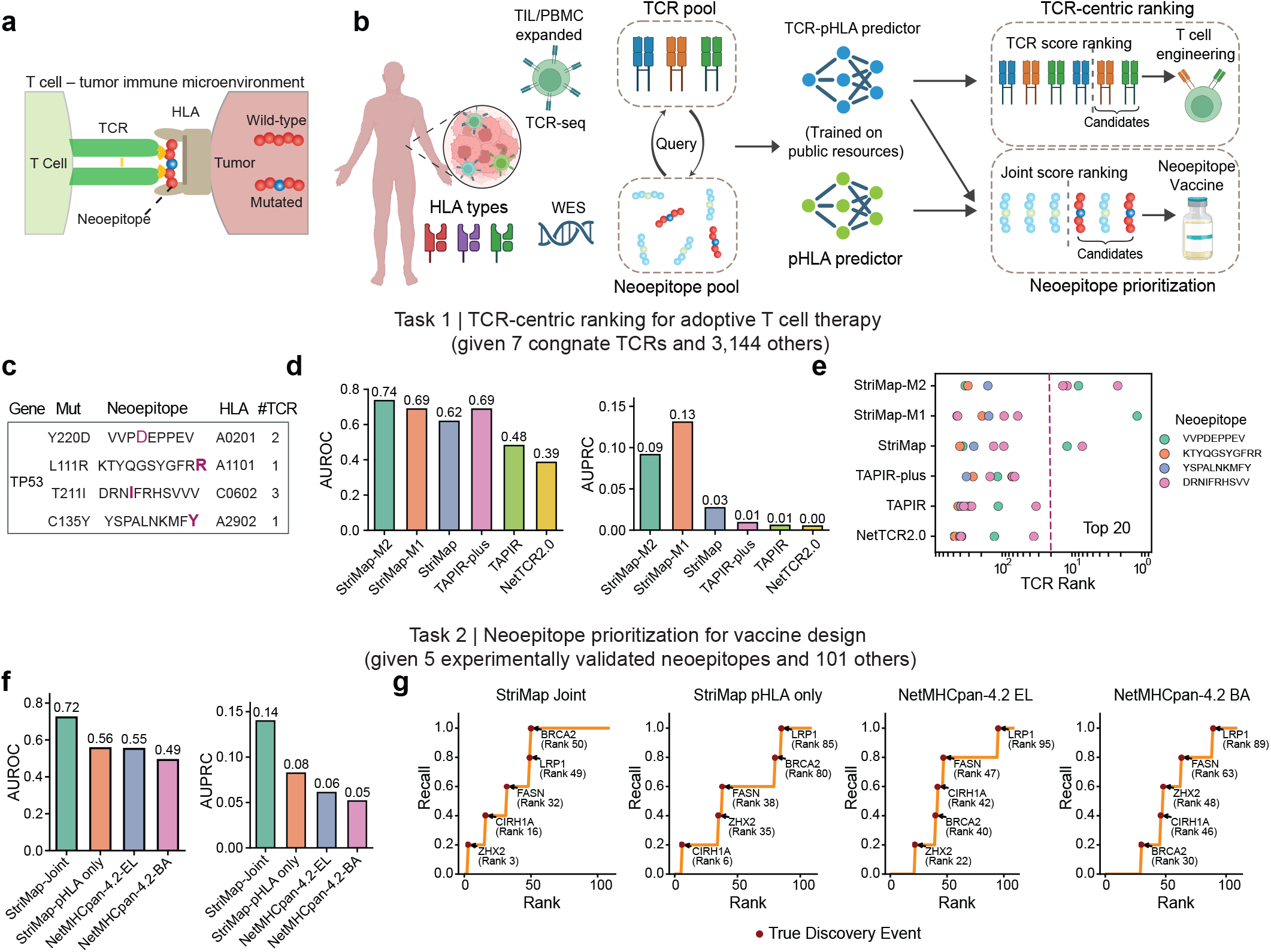
StriMap supports two clinically relevant tasks for T cell-based immunotherapy. StriMap enables two clinically relevant use cases by jointly modeling peptide-HLA presentation and TCR recognition: (i) TCR-centric prioritization for recurrent mutant peptide-HLA pairs in the context of adoptive T cell therapy, and (ii) neoepitope prioritization for personalized cancer vaccine design**(a)** Schematic illustration of T cell recognition within the tumor immune microenvironment, highlighting interactions between TCRs and neoepitope-HLA complexes presented on tumor cells.**(b)** Overview of the StriMap framework for joint analysis of HLA presentation and TCR recognition. Whole-exome sequencing (WES), HLA typing, and TCR sequencing define candidate neoepitopes and patient-specific TCR pools, which are evaluated using StriMap pHLA and TCR-pHLA predictors for downstream prioritization.**(c)** Representative TP53 cancer mutations, their corresponding neoepitopes, and the number of experimentally observed TCRs recognizing each neoepitope.**(d)** Performance comparison of StriMap with TAPIR and NetTCR for prioritizing cognate TCRs given mutant pHLA queries, evaluated by AUROC and AUPRC. TAPIR-plus denotes TAPIR trained on both public datasets and additional in-house proprietary TCR functional data, which StriMap does not access. StriMap-M2 indicates fine-tuning using negative samples generated by randomly mutating one to two amino acids in the peptide, whereas StriMap-M1 indicates fine-tuning using negative samples generated by randomly mutating one to three amino acids in the TCR CDR3α and/or CDR3β regions. For reference, the random baseline is AUROC = 0.5 and AUPRC = 0.002.**(e)** TCR ranking results for TP53-derived neoepitopes. Each point represents a TCR-pHLA interaction for a given neoepitope. Colors denote different neoepitopes, and the dashed line highlights the top 20 ranked TCR candidates.**(f)** AUROC and AUPRC performance of StriMap for prioritizing immunogenic neoepitopes given melanoma patient-specific TCR pools, using a joint score combining TCR-pHLA and pHLA predictors, compared with the pHLA-only StriMap model and NetMHCpan-4.2-EL and NetMHCpan-4.2-BA. For reference, the random baseline is AUROC = 0.5 and AUPRC = 0.047.**(g)** Recall-rank curves illustrating the recovery of true immunogenic neoepitopes across ranking positions for TCR involved predictions versus pHLA predictor only predictions. Red markers denote true discovery events.

We first considered task 1, a TCR-centric setting in which recurrent cancer-associated mutations are known, and the goal is to identify cognate TCRs suitable for adoptive cell therapy. Among these, mutations in TP53 represent one of the most frequent and well-characterized classes of cancer driver alterations across tumor types, and several TP53-derived mutant peptides have been shown to elicit T cell responses in patients^39^. Using representative TP53 hotspot mutations with experimentally observed cognate TCRs originally evaluated by TAPIR^24^ (Fig. 3c), we evaluated StriMap’s ability to rank antigen-specific TCRs for mutant peptide-HLA queries under a fully zero-shot (OOD) setting, where neither the peptide nor the TCRs were seen during training. All positives and negatives were taken directly from TAPIR (7 cognate TCRs; 3,144 negatives), and we compared against TAPIR and the NetTCR-2.0^17^ baseline as reported in the original study^24^.

Within the TCR-centric ranking task, we aim to resolve mutation-specific effects, where mutant peptides differ from their wild-type counterparts by only one or a few amino acids. Distinguishing such subtle differences likely requires amino-acid–level modeling rather than treating non-cognate pairs as the only negatives. To better reflect this biological setting, we introduced a mutation-aware negative sampling strategy, generating negatives via random amino acid substitutions in either the peptide or the TCR CDR3α/β regions. Across both AUROC and AUPRC for TCR ranking, StriMap variants outperformed TAPIR, NetTCR-2.0, and TAPIR-plus (trained with proprietary in-house TCR functional data not accessible to StriMap), with StriMap-M2 (peptide-mutation negatives) achieving the strongest overall performance and StriMap-M1 (TCR-mutation negatives) providing intermediate gains relative to the base model (Fig. 3d). These results indicate that mutation-aware negative sampling is particularly beneficial for modeling cancer-associated antigen recognition.

Consistent with aggregate performance metrics, StriMap showed improved enrichment of true binders among the highest-ranked TCR candidates relative to competing methods (Fig. 3e). For example, multiple experimentally validated TCRs recognizing neoepitopes DRNIFRHSVV and VVPDEPPEV were ranked within the top 20 by StriMap, whereas other methods more often ranked these true positives substantially lower. This capability is critical for T cell therapy, where only a small number of top-ranked candidates can be advanced for costly experimental validation and clinical development. We next asked whether StriMap can resolve mutation-specific effects for a fixed TCR by comparing mutant and wild-type peptide-HLA contexts. Because wild-type peptides may lack immunogenicity either due to limited TCR recognition or low HLA presentation, this comparison helps disentangle the underlying driver. Notably, predicted differences were driven predominantly by changes in pHLA presentation rather than large shifts in inferred TCR–pHLA binding scores (Fig. S14). For example, for the C135Y mutation, the predicted HLA presentation score increased from 0.06 (wild-type) to 0.65 (mutant), whereas the inferred TCR–pHLA score changed only modestly from 0.61 to 0.65, suggesting that the gain in immunogenicity is primarily consistent with enhanced HLA presentation. Together, these results illustrate how jointly modeling pHLA presentation and TCR recognition enables StriMap to predict whether mutation-driven effects are more consistent with altered HLA presentation or altered TCR recognition.

We next evaluated StriMap in task 2, a complementary neoepitope-centric setting, where patient-specific TCR repertoires are available and the objective is to prioritize immunogenic neoepitopes for vaccine development (Fig. 3b). We used paired neoantigen and TCR repertoire data from a melanoma dataset^2^, including clonally expanded TCRs profiled from tumor-infiltrating lymphocytes (TILs) and matched peripheral blood mononuclear cells (PBMCs). In total, this dataset comprised 106 candidate peptide mutations, of which 5 were experimentally confirmed to elicit T cell immune responses. Candidate neoepitopes were ranked using a joint score that integrates predicted neoepitope-HLA presentation with predicted TCR interaction score across the TCR repertoire; we refer to this joint scoring framework as StriMap-joint. Using the mutation-aware StriMap-M2 model, we summarized each candidate neoepitope by the mean interaction score of the top 30 ranked TCR–pHLA pairs, capturing the collective recognition potential of the patient’s expanded T cell repertoire. StriMap-joint consistently achieved higher AUROC and AUPRC than both a pHLA-only StriMap model and NetMHCpan-4.2-based baselines (EL and BA) used in the original study^2^, highlighting the benefit of explicitly incorporating TCR recognition into neoepitope prioritization (Fig. 3f). This advantage was robust to the choice of the number of top-ranked TCR-pHLA pairs used for aggregation (Fig. S15). Recall-rank analysis, summarizing the ranking positions of all experimentally validated immunogenic neoepitopes within the ranked candidate list, further showed that true immunogenic neoepitopes were placed earlier in the ranking when using the joint model compared with pHLA-only or NetMHCpan-based approaches (Fig. 3g).

Together, these results indicate that StriMap’s joint modeling of peptide presentation and TCR recognition can support zero-shot prioritization in two cancer immunotherapy settings. By capturing mutation-driven changes in pHLA presentation and their downstream impact on TCR engagement, StriMap provides a unified framework for TCR selection in adoptive cell therapy and neoepitope ranking for personalized cancer vaccines.

### Proteome-wide discovery of AS-associated bacterial mimic peptides

Having established StriMap’s utility in cancer immunotherapy, we next asked whether the same joint modeling of peptide–HLA presentation and TCR recognition could be extended to autoimmune T cell recognition, where the relevant antigenic triggers are typically unknown. We focused on ankylosing spondylitis (AS), motivated by prior evidence that AS is strongly linked to HLA-B27 and that disease-associated TCRs enriched for a conserved TRBV9 motif may contribute to pathogenesis^40^. Under the molecular mimicry hypothesis, microbial peptides that are presented by HLA-B27 may elicit cross-reactive T cell responses against self-antigens^4^. To define a biologically grounded and clinically relevant search space, we curated a literature-supported cohort of AS-associated gut bacterial strains spanning four major phyla: Bacillota (Firmicutes; including multiple *Streptococcus spp*., *Ruminococcus gnavus*, and others), Verrucomicrobiota (*Akkermansia muciniphila*), Bacteroidetes (*Prevotella/Segatella copri, Prevotella melaninogenica*), and Proteobacteria (e.g., *Klebsiella pneumoniae, Escherichia coli, Parasutterella excrementihominis*)^41–47^. The final panel comprised 16 strains (Table S4), totaling 43,241 annotated proteins used for downstream peptide enumeration.

To evaluate whether StriMap can generalize across patients in a realistic translational regime, we designed a cross-patient few-shot inference workflow centered on TRBV9 TCRs (Fig. 4a). Specifically, we used TCR binding data from patient 1 (TRBV9 TCR AS3.1) and patient 2 (TRBV9 TCRs AS4.1–AS4.4) as reference inputs and then predicted peptide-binding specificity for a query TRBV9 TCR (AS8.4) from an independent patient 3. AS8.4 was selected because it was not among the five TRBV9 TCRs used for few-shot training^40^, enabling an out-of-sample evaluation, and because it showed the strongest activation to the positive-control peptides among the remaining TRBV9 TCRs tested in the original study^40^. This design simulates deployment on a new patient with limited disease-specific supervision while still leveraging large public training resources.

**Fig. 4.**
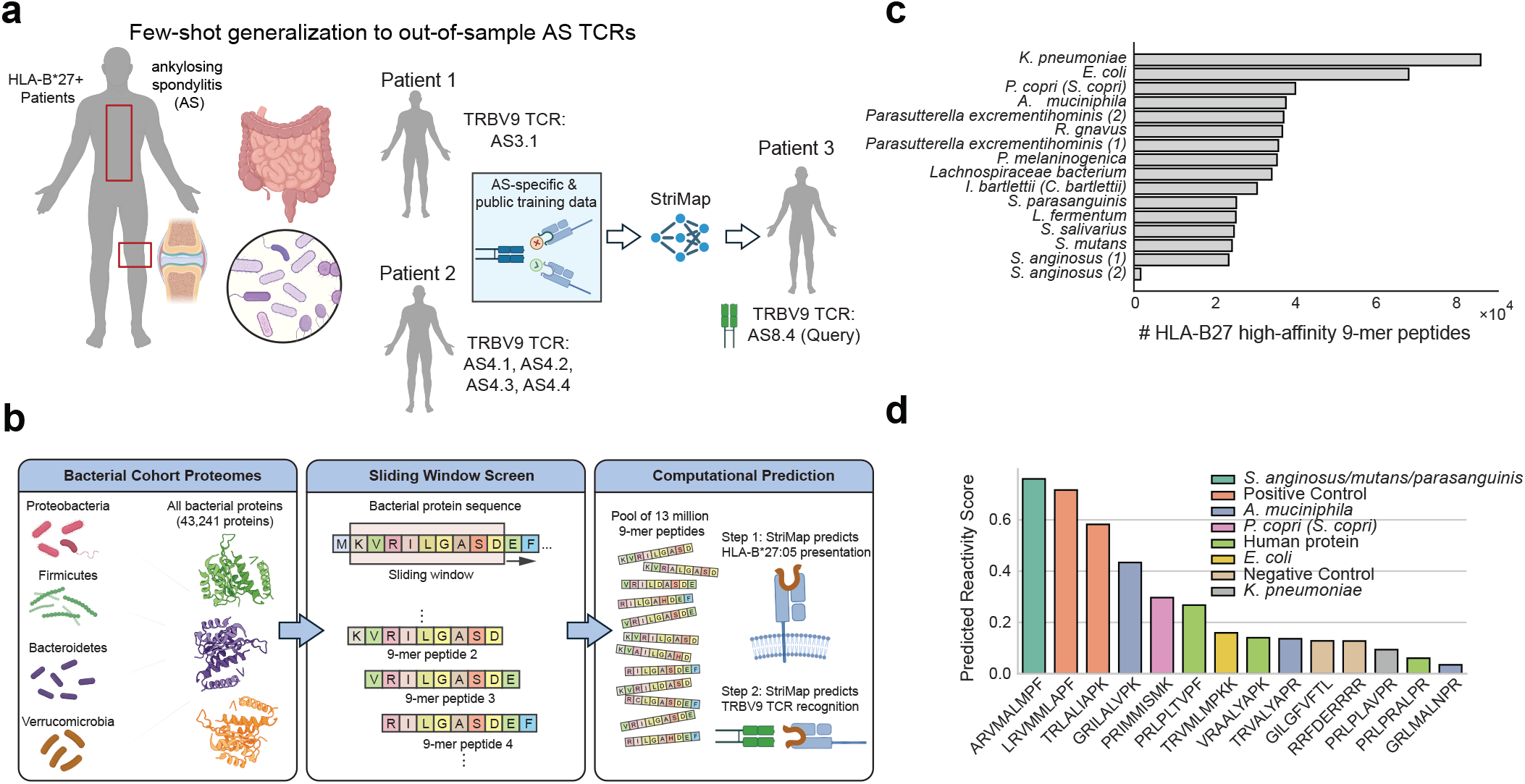
Computational discovery of AS-associated bacterial mimicry peptides through cross-patient prediction. **(a)** AS-associated bacterial cohort and cross-patient few-shot prediction setup. We curated a literature-derived cohort of gut bacterial strains previously reported to be associated with ankylosing spondylitis (AS), spanning four major phyla: Bacillota, including *Streptococcus* spp., *R. gnavus*, etc.), Verrucomicrobiota (*A. muciniphila*), Bacteroidetes (*P. copri, P. melaninogenica*), and Proteobacteria (*K. pneumoniae, E. coli, Parasutterella excrementihominis*). To assess generalization to unseen patient TCRs, we implemented a cross-patient inference workflow in which TRBV9 TCR binding data from Patient 1 (AS3.1) and Patient 2 (AS4.1–AS4.4) were used as reference inputs to train the specificity model, and StriMap then predicted peptide-binding specificity for a query TRBV9 TCR from an independent Patient 3 (AS8.4), simulating a few-shot setting.**(b)** Large-scale in silico 9-mer peptide screening pipeline. The proteomes of the selected bacterial cohort, comprising 43,241 proteins, were retrieved and processed. A sliding window approach with a window size of k=9 and a stride of 1 amino acid was applied to generate a pool of approximately 13 million 9-mer peptides. The StriMap computational framework performs a dual-task prediction: Task 1 predicts the likelihood of HLA-B*27:05 presentation, and Task 2 predicts the recognition probability by specific TRBV9 TCRs (e.g., AS8.4).**(c)** Landscape of HLA-B27 high-affinity 9-mer peptides across the cohort. The bar chart illustrates the distribution and total count of predicted high-affinity 9-mer peptides for each bacterial strain. Strains such as *K. pneumoniae* and *E. coli* exhibit a high density of potential HLA-B27 binders, highlighting their significance in the molecular mimicry hypothesis.**(d)** Predicted reactivity scores for candidate bacterial peptides. Representative peptides from various phyla were ranked by their predicted reactivity scores against the target TCR. High-scoring candidates from *Streptococcus spp*. and *A. muciniphila* are compared against positive and negative controls, indicating potential cross-reactivity between bacterial antigens and AS-associated TCRs.

We then performed proteome-wide in silico screening of bacterial peptides at the scale required for mimicry discovery (Fig. 4b). For each bacterial proteome, we generated overlapping 9-mer peptides using a sliding window (k = 9, stride = 1), yielding a pool of 13,068,902 candidate 9-mers. StriMap was then applied in a two-stage, dual-task manner: (*i*) predicting the score of HLA-B*27:05 presentation, followed by (*ii*) predicting recognition by the disease-associated TRBV9 TCRs, thereby coupling presentation and recognition in a single screening workflow. In our implementation, we retained the top predicted HLA-B*27:05 binders (top 3% relative to a random-peptide background; 568,982 peptides) for downstream TCR screening (Fig. 4c). After filtering for high-confidence HLA-B*27:05 candidates, we used the StriMap TCR–pHLA predictor to estimate interactions between each candidate peptide and five disease-associated TRBV9 TCRs (AS3.1, AS4.1–AS4.4) previously reported to be reactive to multiple HLA-B*27:05–bound peptides^40^. Peptides were then ranked by the mean predicted binding score across the five TCRs, and we selected seven top-ranking bacterial peptides that were each predicted to interact strongly with at least two TRBV9 TCRs. We imposed this multi-TCR criterion to reflect the known cross-reactivity of disease-associated TRBV9 TCRs and to prioritize candidates with broader potential to activate multiple TRBV9 clonotypes instead of peptides specific to a single TCR, thereby increasing translational relevance. We next evaluated the predicted reactivity of these candidates against the out-of-sample query TRBV9 TCR AS8.4, and benchmarked candidate bacterial peptides against positive and negative controls as well as human-derived peptides included as specificity checks. Consistent with the mimicry hypothesis, top-ranked candidates included peptides derived from *Streptococcus spp*. and *A. muciniphila*, and multiple high-scoring candidates (e.g., ARVMALMPF, GRILALVPK, PRIMMISMK, PRLPLTVPF) emerged among the leading predictions under this cross-patient few-shot setting (Fig. 4d).

### Experimental validation of TRBV9^+^T cell activation by bacterial peptides

To experimentally validate StriMap-prioritized candidate mimic peptides in an HLA-B*27:05– restricted setting, we implemented a discovery workflow in which computational screening nominates candidate TCR–peptide–HLA interactions for targeted testing and iterative refinement (Fig. 5a). We generated endogenous TCR-knockout Jurkat cell lines and re-expressed the TRBV9 TCR AS8.4 by lentiviral transduction (Materials and Methods). Antigen-presenting cells (APCs) were generated by transducing 293T cells to express HLA-B*27:05. T cells were then co-cultured with APCs that had been pre-incubated with the candidate peptides, and T cell activation was assessed by measuring both CD69 surface expression and fluorescence from the NFAT-driven ZsGreen reporter, in which ZsGreen expression is controlled by a nuclear factor of activated T cells (NFAT) response element to report downstream TCR signaling (Fig. 5b; Materials and Methods).

**Fig. 5.**
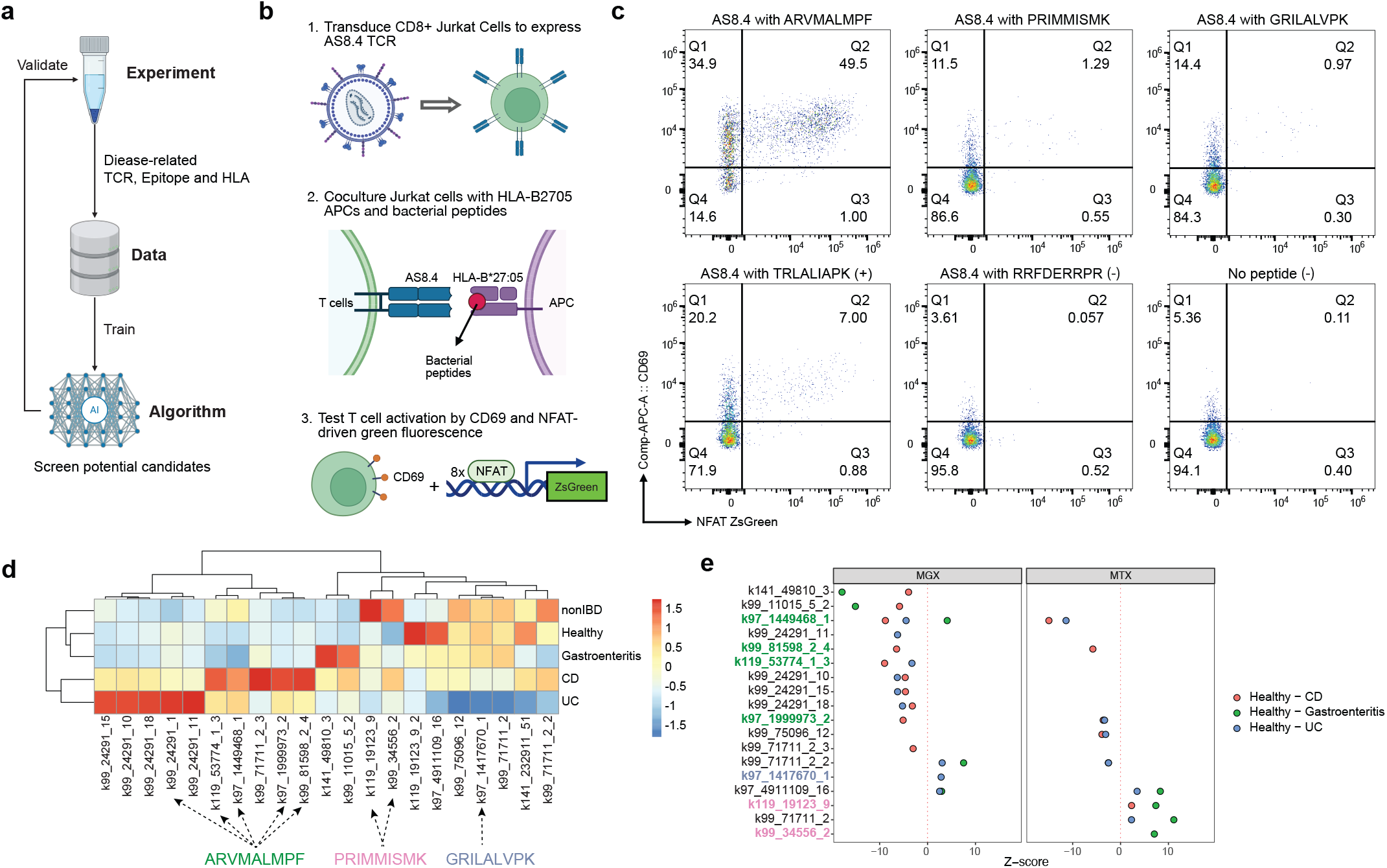
Experimental validation of AS-associated microbial mimicry peptides. **(a)** Overview of discovery framework, in which StriMap prioritizes candidate disease-associated TCR-peptide-HLA interactions for experimental validation and iterative refinement.**(b)** Experimental workflow for validating interactions between the AS-associated TCR AS8.4 and candidate peptides. CD8^+^ Jurkat T cells were transduced to express the AS8.4 TCR and co-cultured with HLA-B*27:05-expressing antigen-presenting cells (APCs) pulsed with candidate bacterial peptides.**(c)** Flow cytometry analysis of T cell activation following peptide stimulation. Activation was quantified by co-expression of CD69 (y-axis) and NFAT-driven ZsGreen reporter activity (x-axis). Three bacterial peptides induced robust T cell activation, comparable to the positive control, whereas negative and background controls showed minimal activation. Activated cells are enriched in the CD69^+^ZsGreen^+^ quadrant.**(d)** Metagenomic and metatranscriptomic profiling of bacterial peptide candidates across clinical cohorts. Heatmap shows the relative prevalence of peptide-associated features in healthy controls, infectious gastroenteritis, Crohn’s disease (CD), and ulcerative colitis (UC), following Z-score normalization.**(e)** Statistical comparison of peptide-associated features across diagnostic groups. A Kruskal-Wallis test was performed across healthy, infectious gastroenteritis, UC, and CD cohorts. Features with significant group-level differences (p < 0.05) were further assessed using post hoc Dunn’s tests with multiple-comparison correction.

Although not all candidates were predicted to bind AS8.4, we tested the full set of 10 peptides to directly assess the model’s ability to separate activators (CD69^+^ZsGreen^+^) from non-binders. We observed robust T cell activation in response to the three predicted positive binders: ARVMALMPF (from *Streptococcus anginosus, S. mutans*, and *S. parasanguinis*; 49.5%), PRIMMISMK (from *Prevotella copri* and *Segatella copri*; 1.29%), and GRILALVPK (from *Akkermansia muciniphila*; 0.97%) (Fig. 5c; Fig. S16). For comparison, we included two previously reported positive controls (TRLALIAPK, 7.0%; LRVMMLAPF, 57.8%), and two negative controls (RRFDERRPR, 0.057%; GILGFVFTL, 0.084%) (Fig. 5c; Fig. S16), together with background conditions (no peptide; no APC/no peptide), all of which showed minimal activation (Fig. 5c; Fig. S16).

StriMap predictions across the ten peptides were highly concordant with experimental outcomes: the three activating peptides (ARVMALMPF, GRILALVPK, PRIMMISMK) were ranked among the top candidates, whereas non-activating peptides received consistently low scores. To contextualize these results, we benchmarked multiple modeling strategies on the experimentally tested peptide–TCR pairs by labeling the three activating peptides as positives and the remaining peptides as negatives, and computing AUROC/AUPRC from each method’s predicted scores (Fig. S17). StriMap substantially outperformed logistic regression, the amino-acid frequency– based baselines used in Yang et al. (2022)^40^, as well as NetTCR-2.2^18^, in both metrics. Notably, incorporating limited experimental feedback in a few-shot setting improved performance, indicating that StriMap can efficiently leverage sparse experimental validation data to refine disease-specific TCR-antigen recognition predictions.

Finally, given reports linking potentially pathogenic *Streptococcus* species to AS, IBD, and AS– IBD overlap^41,45^, we asked whether the experimentally validated candidates show broader clinical associations. We performed an exact k-mer search against a translated, non-redundant microbial protein catalog constructed from metagenomic mix-assemblies^48^. This protein catalog was built from 156 individuals with IBD including ulcerative colitis (UC, n = 29) and Crohn’s disease (CD, n = 82) as well as from healthy controls (n = 45) across IBD cohorts^49–51^, healthy controls from the Men’s Lifestyle Validation Study (MLVS) study (n = 361)^52^, and 970 patients with pathogen-confirmed gastroenteritis^53^. Matched metatranscriptomic data were available for all metagenomic samples, allowing us to evaluate the transcriptional activity of the genes encoding proteins that contain the candidate peptides. Across the ten candidate peptides, five proteins containing the ARVMALMPF motif were significantly more prevalent and significantly enriched in patients with CD or UC compared to healthy controls, whereas two proteins with PRIMMISMK were more prevalent and transcriptionally upregulated in non-IBD samples (Fig. 5d, e; Table S5). In particular, ARVMALMPF—predicted with the highest score in the screening stage (Fig. 4d) and validated here to activate AS8.4^+^ T cells—showed consistent enrichment in IBD cohorts, supporting the possibility of shared microbial triggers between AS and IBD.

## Discussion

Deep learning has enabled major progress across biomedicine, yet generalizing to new biological contexts remains challenging, particularly under data sparsity, class imbalance, and dataset bias. Rather than relying solely on ever-larger datasets, we focus on developing models that better capture biological structure under limited supervision, and apply this principle to TCR antigen-specificity prediction, a central problem in infection, cancer, and autoimmunity. T cell–mediated immunity relies on a recognition cascade in which peptides are presented by HLA molecules and subsequently engaged by diverse T cell receptors. Modeling this tri-molecular process has been difficult because the interaction space is combinatorially large, recognition is structurally constrained yet flexible, and available experimental data are sparse and biased. Here we present StriMap, a unified framework that jointly models peptide–HLA presentation and TCR recognition using complementary physicochemical, sequence-context, and structure-informed representations.

A central conceptual advance of StriMap is its treatment of antigen presentation and TCR recognition as a coupled biological process rather than independent prediction tasks. By conditioning TCR recognition on the peptide–HLA landscape, StriMap provides a biologically consistent decision pathway that helps disentangle whether changes in immunogenicity arise from altered presentation or altered TCR engagement. Moreover, StriMap incorporates structure-derived constraints at scale without requiring experimentally solved complexes, enabling reasoning about interface geometry beyond current structural databases. In practice, we find that training strategy is a critical complement to architecture: dynamic and mutation-aware negative sampling exposes the model to realistic near-miss decoys, which is particularly important in clinical settings where neoantigens often differ from self by a single amino acid.

StriMap’s utility is highlighted by applications in two distinct disease settings where candidate selection must be aggressively prioritized. In cancer immunotherapy, StriMap improves enrichment of true binders among the top-ranked candidates under zero-shot conditions, a setting that mirrors clinical reality because patient-specific neoantigens are inherently novel. The coupled modeling of presentation and recognition provides a practical framework for prioritizing TCRs and vaccine neoepitopes when only a small number of candidates can be experimentally tested. Although we focus on melanoma and TP53 mutations here, we anticipate that this prioritization strategy may generalize to other cancer contexts. In autoimmunity, StriMap links a large-scale microbial sequence space to antigen-specific T cell responses: by combining disease-associated TRBV9 TCR clonotypes with an HLA-B*27:05-restricted microbial peptide landscape, we identified bacterial peptides that were experimentally validated to activate AS-associated T cells. The enrichment of one validated peptide in IBD cohorts further suggests that shared microbial antigenic drivers may contribute to overlapping inflammatory phenotypes, illustrating how StriMap can convert large-scale in silico screening into testable mechanistic hypotheses.

Several limitations and opportunities remain. First, because TCR–pHLA data are sparse and unevenly distributed across epitope space, fully zero-shot generalization to entirely novel epitopes remains challenging. Our results therefore suggest that evaluation of zero-shot performance should emphasize functional prioritization, rather than relying solely on aggregate metrics such as AUROC. OOD benchmarks such as IMMREP23 include only a small number of unseen epitopes (n = 3), limiting the interpretability of global performance statistics. In contrast, StriMap demonstrates strong practical utility in clinically relevant ranking settings: it placed validated TP53-specific TCRs among the top-ranked candidates in cancer immunotherapy, and prioritized a small set of activating peptides from a pool of ∼13 million bacterial 9-mers in autoimmunity. Such top-K precision is often the key determinant of translational success, because it directly governs which candidates are selected for costly experimental validation. Nevertheless, few-shot adaptation using limited task- or patient-specific data remains a reliable strategy for further improving accuracy in applications requiring higher fidelity.

Second, structure-informed representations improve interface modeling but may not capture conformational dynamics that influence recognition; incorporating structural uncertainty (e.g., ensemble predictions) or dynamics-inspired features may further improve predictions. Consistent with this direction, the ongoing IMMREP25 Kaggle challenge underscores the importance of structure-aware TCR–pHLA modeling, although direct comparison is currently limited because top-performing structure-based approaches have not yet been published. Moreover, many leading models are based on AlphaFold3^54^ prediction, whose computational cost and throughput constraints limit applicability to proteome-scale screening, whereas StriMap leverages ESMFold for efficient large-scale prioritization. Third, our current implementation focuses on HLA class I and CD8^+^ T cell recognition; extending the framework to HLA class II presentation and CD4^+^ T cell recognition will be important for broadening translational impact.

To facilitate broad adoption and iterative improvement, we provide a web portal and training interface that support both prediction and rapid fine-tuning on local datasets, enabling users to incorporate sparse experimental feedback while retaining knowledge from large-scale pretraining (Fig. S18). We anticipate that this combination of coupled modeling, structure-aware representations, and lab-in-the-loop validation will help bridge computational prediction and mechanistic immunology, enabling more effective discovery in cancer immunotherapy and antigen-driven autoimmune disease.

## Methods

### The StriMap Framework

StriMap is a unified deep learning framework for modeling pHLA binding and TCR recognition of pHLA complexes at residue resolution. The framework consists of two modular components: (*i*) a peptide-HLA predictor that learns transferable pHLA representations, and (*ii*) a TCR-pHLA predictor that leverages pretrained pHLA features. Both components are built upon a unified Sequence and Structure Feature Extractor (SSFE) and attention-based residue-level interaction modeling using Bilinear Attention Networks (BAN).

### Sequence and Structure Feature Extractor (SSFE)

Given an input protein sequence *S* of length *L*, SSFE constructs residue-level representations by integrating physicochemical, sequence-context, and structure-informed features. All three modalities are encoded independently and subsequently fused.

#### Physicochemical Encoding

Each residue <u>*s*_*i*_</u> is encoded using a 20-dim physicochemical descriptor vector 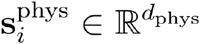 derived from AAindex^27^ (selected properties, such as hydrophobicity, charge, polarity, molecular volume, and side-chain flexibility; z-score normalized across amino acids) (Table S1). For a sequence of length <u>*L*</u>, physicochemical features are assembled into a matrix

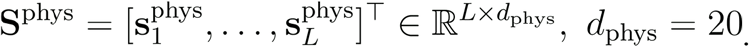

To align this modality with the model’s latent space, we apply a fully connected layer to map the features to *d* dimensions:

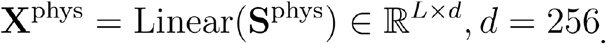

#### Sequence Context Encoding

To capture evolutionary and contextual information beyond local amino acid properties, we employ a pretrained protein language model, ESM2 (650 million parameters)^28^. For an input sequence *<u>S</u>*, residue-level semantic embeddings are extracted from the final transformer layer of ESM2 followed by a fully connected layer:

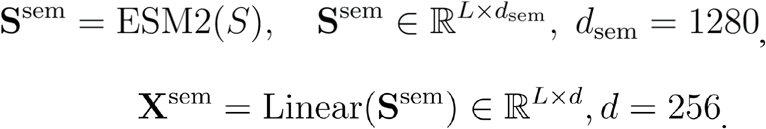

#### Structure Geometric Encoding

Protein structural information is derived from 3D structures predicted by ESMFold. For each residue <u>*s*_*i*_</u>, we extract a structural descriptor vector 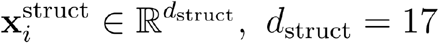, which includes backbone torsion angles (encoded as sine and cosine pairs), secondary structure states, relative solvent accessibility, residue contact counts, and predicted confidence scores (pLDDT). In addition, the Cartesian coordinates of the Cα atom are retained:***r***_*i*_ ∈ ℝ^3^.

To explicitly model spatial relationships, residues are represented as nodes in a geometric graph, and a stacked SE(3)-equivariant graph neural network (EGNN) is applied. For EGNN layer *<u>ℓ</u>*, node features and coordinates are updated as:

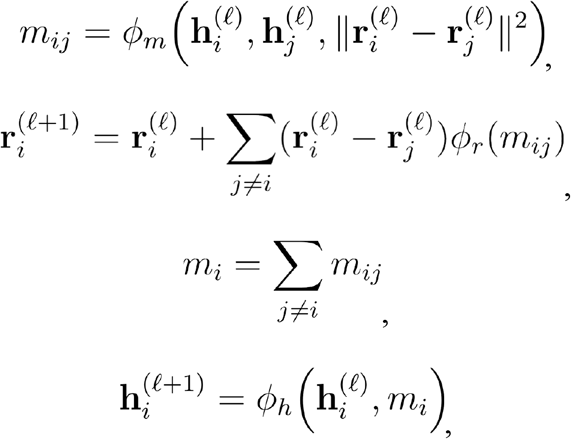

where *ϕ*_*m*_, *ϕ*_*r*_, and *ϕ*_*h*_ are learnable multilayer perceptrons. Node features are initialized as 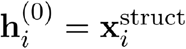 and 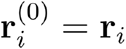. The coordinate update formulation guarantees SE(3)-equivariance.

Finally, the structure-derived geometric encoding is defined as the node embeddings from the final layer followed by a fully connected layer:

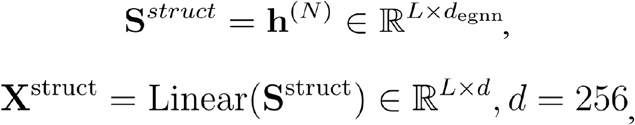

where *d*_*egnn*_ denotes the hidden dimension of the EGNN.

#### Multimodal Feature Fusion

Physicochemical and sequence-context features are fused using a gated cross-attention mechanism. Given **X**^phys^ and **X**^sem^, cross-attention is computed as:

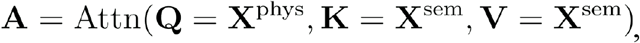

where

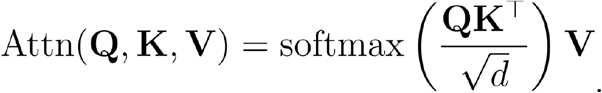

A learnable gating function controls modality contribution:

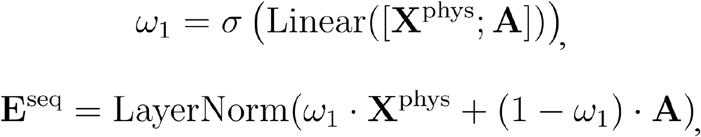

Structure-aware embeddings are further integrated with sequence-derived features. The final residue representation is computed as:

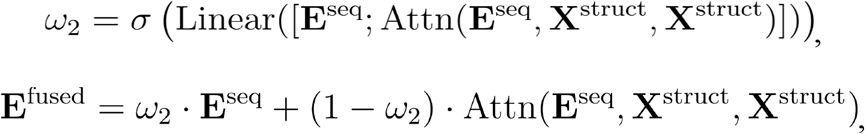

where ω_*k*_ ∈ ℝ, *k* ∈ {1,2} is a scalar gating coefficient obtained via a sigmoid activation function . σ( ·)

### Interaction Modeling: Bilinear Attention Networks (BAN)

To model the fine-grained contact interface and capture the intricate dependencies between the residues of two protein chains (e.g., Peptide and HLA, or TCR and pHLA), we implement Bilinear Attention Networks. Unlike standard dot-product attention which often reduces information into a single dimension, BAN models pairwise residue interactions via a bilinear compatibility function and aggregates interaction-aware features without explicitly constructing a full outer-product tensor.

We utilize learnable weight matrices **U** ∈ ℝ^D×D′^ and **V** ∈ ℝ^D×D′^ to project two representations 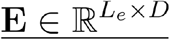 and 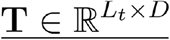 into a joint space. The transformed features for the *i*-th residue of **<u>E</u>** and the *j*-th residue of **<u>T</u>** are:

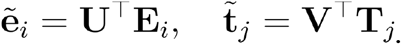

The interaction score <u>**S**_*i,j*_</u> for each residue pair is computed using a learnable parameter **q** ∈ ℝ^D′^ to measure bilinear compatibility as:

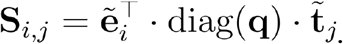

The attention map **I**_*i,j*_ is normalized via an explicit 2D Softmax function:

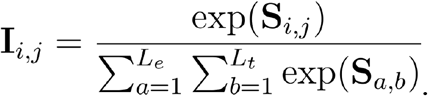

The pooled interaction vector **z** is derived through the Hadamard product (⊙) across all residue pairs:

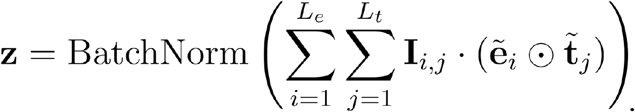

### Peptide-HLA Predictor Model

The peptide-HLA predictor integrates SSFE, Transformer encoders, cross-attention, and BAN. Let *S*^pep^ and *S*^hla^ denote the amino acid sequence of peptide and HLA. Peptide and HLA residue features are first encoded independently by SSFE module:

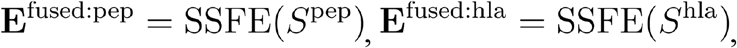

and Transformer blocks:

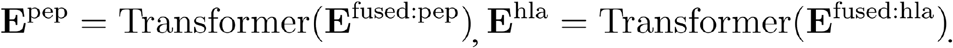

Bidirectional cross-attention is then applied:

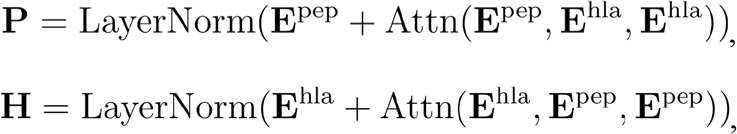

BAN is applied between peptide residues and HLA pocket residues predefined as 34 canonical binding-groove positions^10^, producing an interaction vector,

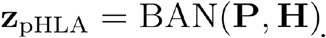

which is passed to a multilayer perceptron with a sigmoid activation to predict peptide-HLA binding score.

### TCR-Peptide-HLA Predictor Model

The TCR-pHLA predictor model predicts the specific recognition of a pHLA complex by a TCR α/β heterodimer. This model utilizes a multi-stage training pipeline and a sophisticated interaction hierarchy to handle the increased complexity of the quaternary complex.

#### Feature Aggregator: Pre-training and Transfer Learning

To stabilize and guide the learning of TCR-pHLA recognition, the pHLA predictor is first pre-trained and subsequently reused as a feature aggregator. This transfer learning strategy ensures that the model leverages established knowledge of pHLA complex stability before modeling the TCR interaction.

Let **H**^pre^ and **P**^pre^ denote the pre-trained embeddings (for HLA and peptide, respectively) extracted from the frozen pHLA predictor, and let **H**^task^ and **P**^task^ denote the task-specific embeddings generated within the TCR-pHLA pipeline. These representations are integrated through gated fusion:

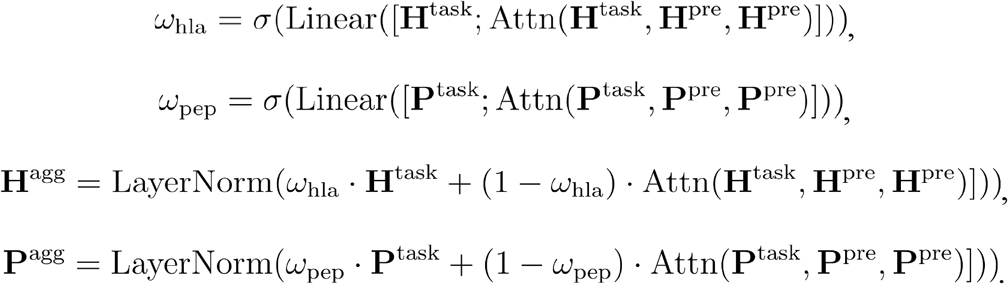

This aggregation mechanism allows the TCR model to adaptively focus on pHLA binding features that are most relevant to TCR recognition while maintaining a robust foundation of binding stability.

#### Interaction Hierarchy: SSFE + Transformer + Cross Attention + BAN

Following feature aggregation, the TCR-pHLA interaction is modeled through a hierarchical sequence of operations:

- Let *S*^tcra^ and *S*^tcrb^ denote the amino acid sequences of TCRα and TCRβ chains. The residue features are encoded independently by SSFE module and Transformer blocks:**E**^tcra^ = Transformer (SSFE) (S^tcra^)), **E**^tcrb^ = Transformer (SSFE) (*S*^tcrb^)).
- CDR3 Segment Extraction: The model identifies and extracts the hypervariable CDR3α and CDR3β segments. For each chain, let 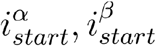 and 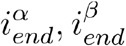 be the anchor residue indices. The segment **E**^cdr3a^ and **E**^cdr3a^ is extracted as: 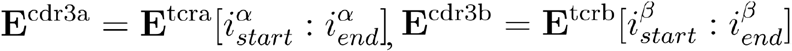
- Multichannel Cross-Attention: Mutual information is exchanged between the TCR loops and the pHLA aggregator features to refine the docking orientation:

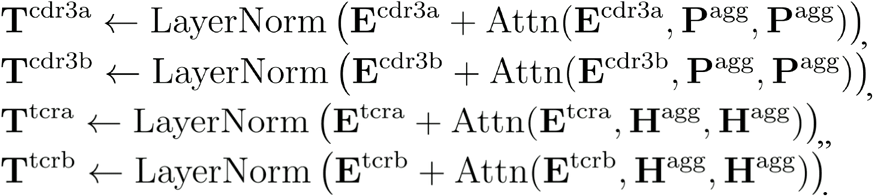
- Bilinear Interaction Modeling: Using the BAN architecture described in Section 3.3, the model computes four distinct interaction vectors:

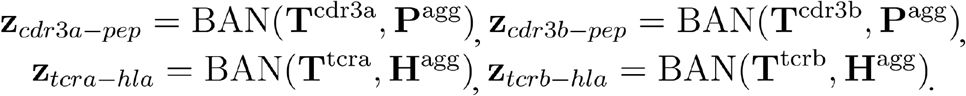
- Prediction Head: These vectors are concatenated into a global interaction representation **z**_*total*_ and passed through a multi-layer perceptron (MLP) with a sigmoid activation to output the final TCR-pHLA binding score.

### Negative Sampling

To improve discrimination and mitigate shortcut learning under limited experimentally validated negatives, we used two complementary negative-sampling strategies: dynamic shuffling-based negatives and mutation-aware hard negatives.

#### Dynamic Negative Sampling

Because comprehensive negative binding data are scarce for TCR-pHLA interaction prediction, we generated negative pairs for training. Common strategies include: (i) fixing pHLA and randomly shuffling TCRs to form mismatched pairs, while excluding any known positive pairs; and (ii) pairing pHLA with background TCRs from healthy donors. We adopted the shuffling-based strategy to avoid potential distribution mismatches between positive and negative TCR repertoires that may arise when using healthy-donor backgrounds^35^. Concretely, we sampled negatives at a 3:1 negative-to-positive ratio. Rather than fixing a single set of negatives before training, we regenerated shuffled negatives dynamically during training (e.g., each epoch), which expands the effective negative space while keeping the positive set unchanged and reduces reliance on a limited, static negative pool.

#### Mutation-aware hard negative sampling

In the cancer mutation analyses, we additionally introduced a mutation-aware strategy to construct hard negatives that more closely reflect the biological challenge of distinguishing mutants from closely related self-contexts. Specifically, we generated decoy pairs via small sequence perturbations:

- **Peptide mutation**: for a positive peptide, we randomly substitute 1-2 residues to create minimally perturbed decoys that are expected to reduce peptide-HLA and/or TCR recognition compatibility.
- **TCR CDR3 mutation**: we randomly mutate 1-3 residues within the CDR3 loop(s) (excluding conserved anchor positions) to produce near-neighbor decoys likely to alter specificity.

We note that this strategy can introduce label noise, as a subset of perturbed sequences may still retain measurable binding. Nevertheless, under this construction, the experimentally supported positives are expected to exhibit higher binding propensity on average than the mutated decoys, making these hard negatives a useful approximation for training and evaluation in the absence of exhaustive negative measurements.

### Training Objective

Models are trained using focal loss to address class imbalance. For label *y* ∈ {0,1} and predicted probability p, the loss is defined as:

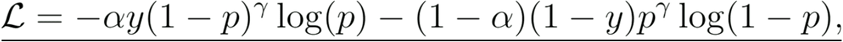

where <u>α</u>balances positive and negative classes and <u>γ</u>emphasizes hard examples. This strategy forces the model to distinguish true molecular recognition from highly similar decoy interactions, substantially improving specificity.

### Training Details

- Optimizer: AdamW with a learning rate of 1 × 10 ^− 4^ and weight decay.
- Regularization: Dropout of 0.1 and Gradient Clipping at a max norm of 2.0.
- Cross-Validation: 5-fold cross-validation with an EarlyStopping patience of 5 epochs for Peptide-HLA predictor, and 10-fold cross-validation with an EarlyStopping patience of 10 epochs for TCR-pHLA predictor, both based on validation AUROC. The maximum epoch was set to 100.
- Batch size: The batch size was set to 256.
- Hardware: All models were trained on NVIDIA A6000 GPUs using the PyTorch framework.

### Experimental Details

#### Plasmid construction

The AS8.4 TCR sequences TRBV901-TRBJ2-301 and TRAV2101-TRAJ2801 were obtained from the international ImMunoGeneTics information system (IMGT/GENE-DB, https://www.imgt.org/). CDR3 regions correspond to those reported by Yang et al. (2022) (β: CASSVGTYSTDTQYF; α: CAVNSPGSGAGSYQLTF). Gene fragments were codon-optimized for Homo sapiens, joined with a P2A linker, and assembled into a pLX307 vector. Correct assembly was confirmed by Sanger sequencing (CCIB DNA Core, Massachusetts General Hospital, Boston).

#### Cell lines

Jurkat E6.1 (ATCC; TIB-152), HEK293T (ATCC; CRL-3216), and Plat-A (Cell Biolabs, San Diego, CA, USA; RV-102) cells were cultured at 37 °C and 5 % CO_2_. Jurkat and Plat-A cells were maintained in RPMI-1640 (Thermo Fisher; 61870036); HEK293T and Plat-A cells were maintained in DMEM (Thermo Fisher; 10569010). Media were supplemented with 10 % (v/v) heat-inactivated FBS (Thermo Fisher; 26140079) and 100 µg ml^−1^ penicillin-streptomycin (Thermo Fisher; 15140-122). Cell lines were screened mycoplasma-free every 3 months with the Mycoplasma Detection Kit (InvivoGen; rep-mys-50).

#### CRISPR-Cas9 knockout of endogenous TCR

To eliminate endogenous TCR expression in Jurkat cells, both the TCR α constant region (TRAC) and the TCR β constant region (TRBC1) were targeted using synthetic guide RNAs (sgRNAs) complexed with Cas9-GFP protein (IDT; 10008100). Ribonucleoprotein complexes were assembled at a 1:2 molar ratio of Cas9-GFP (IDT; 10008100) to sgRNA (IDT). The following sgRNA sequences were used: for TRAC, 5′-TGGATTTAGAGTCTCTCAGC-3′ and 5′-ACAAAACTGTGCTAGACATG-3′; for TRBC1, 5′-CGTAGAACTGGACTTGACAG-3′ and 5′-CCCACCAGCTCAGCTCCACG-3′. Jurkat cells were electroporated with ribonucleoprotein complexes using a 4D-Nucleofector™ X Unit (Lonza) and the SE Cell Line 4D-Nucleofector™ X Kit S (Lonza; V4XC-1032). Program CL-120 was used, and cells were transferred immediately to 2 mL pre-warmed RPMI + 10 % FBS. After 5 days, cells were stained with PE-conjugated anti-human TCRαβ (BioLegend; 306708). TCRαβ^−^ cells were sorted with a Bigfoot™ Spectral Cell Sorter (Thermo Fisher Scientific).

#### Lentiviral vector production and transduction

HEK293T cells were co-transfected with the plasmids psPAX2 (Addgene #12260) and pMD2.G (Addgene #12259) at 4:4:1 ratio using TransIT®-LT1 (Mirus Bio; MIR 2300). Supernatants were harvested after 72 hours, filtered, and stored at −80 °C. Viral concentrations were determined with Lenti-X™ GoStix™ Plus (Takara Bio, 631280). TCR knockout Jurkat cells were transduced with the virus by spinfection (1,000 × g, 90 min, 32 °C) in the presence of 8 µg/mL polybrene (Sigma-Aldrich; TR-1003-G). After 48 h, 2 µg/mL puromycin was applied for 5 d. Expression of AS8.4 TCR was verified by staining with PE-conjugated anti-human TCRαβ (BioLegend; 306708) and APC-conjugated anti-human CD3 (BioLegend; 300411).

#### Retroviral delivery of CD8αβ and NFAT-ZsGreen reporter

The CD8α-CD8β-8×NFAT-ZsGreen construct (Addgene #153417) was packaged in Plat-A cells. TransIT-LT1 (Mirus Bio; MIR 2300) was used for transfection. Supernatants (48 h, 72 h) were filtered and concentrated 10-fold with the Lenti-X Concentrator (Takara Bio; 631231). Jurkat-AS8.4 cells were spin-infected as above, cultured 72 h, and sorted for CD8α^+^.

#### Generation of HEK293T-B2M-HLA-B*27:05 antigen-presenting cells

Lentiviral construct encoding human β_2_-microglobulin (B2M) and HLA-B*27:05 was packaged as above. HEK293T cells were transduced by spinfection (1,000 × g, 90 min, 32 °C) in the presence of 8 µg/mL polybrene, selected with puromycin (2 µg/mL, 5 d), and expression assessed with FITC anti-HLA-B27 (Invitrogen; MAB1285).

#### Peptide stimulation assay

HEK293T-HLA-B*27 cells (1 × 10^5^ cells per well) were seeded in 96-well U-bottom plates in 100 µL DMEM complete medium. After 24 h, synthetic peptides (100 µM final; GenScript) were pre-incubated with HEK293T-HLA-B*27 cells for 60 min at 37 °C and 5 % CO_2_. Then 1 × 10^5^ Jurkat-AS8.4-NFAT-ZsGreen cells in 100 µL RPMI + 10 % FBS were added. Co-cultures proceeded 24 h (37 °C, 5 % CO_2_).

Cells were then harvested and stained with APC anti-human CD69 (BioLegend; 985206) in PBS + 2 % FBS. Data were acquired on Cytek® Aurora (Cytek Biosciences, Fremont, CA) using SpectroFlo v3.2.0. Analysis was performed in FlowJo™ v10.10.0 (BD Biosciences, San Jose, CA, USA). Live singlets (FSC-A vs SSC-A, FSC-H) were gated for ZsGreen and CD69. Activation was expressed as % CD69^+^ and ZsGreen^+^.

## Acknowledgements

We thank Q. Gong, C. Li, F. Chen, C. Krishna, W. Jin, B. Shao, L. Yuan, X. Zhang, J. Zhang, R. Zhang, and S. Liu for valuable discussions regarding method design. We thank E. Forte and T. Reimels for helpful discussions and feedback on the manuscript. Figures 1, 3, 4 and 5 were partially created with BioRender.com

## Code availability

The open-source StriMap codebase is maintained at https://github.com/uhlerlab/strimap-tools. A publicly accessible web server implementing the StriMap prediction platform is available at www.strimap.com, providing batch prediction, analysis, visualization and model fine tuning for peptide-HLA binding and TCR-pHLA recognition.

## Data and materials availability

All benchmarking data analyzed in this article are publicly available through original publications. The final training data and AS-related microbial peptides analyzed in this study are provided at https://doi.org/10.5281/zenodo.18002170.

## Author contributions

R.J.X. and C.U. directed the study. K.C., M.S., E.M.B., and P.N.U.N. conceived the study. K.C. developed the model. K.C., M-M.P. and J.P. performed data analysis. D.B.G. and R.L. designed the AS experiments. R.L. performed the AS experiments. O.A. and D.B.G provided crucial insights. K.C., R.L., M.S., M-M.P., O.A., D.B.G, C.U. and R.J.X. wrote the manuscript.

