## Supplementary tables and figures for "A structure-informed deep learning framework for modeling TCR-peptide-HLA interactions"

| AAindex1 ID | Property (description) | Reference |
|---|---|---|
| BIGC670101 | Residue volume | Bigelow (1967) |
| CHAM820101 | Polarizability parameter | Charton–Charton (1982) |
| CHOP780201 | Normalized frequency of alpha-helix | Chou–Fasman (1978b) |
| CHOP780202 | Normalized frequency of beta-sheet | Chou–Fasman (1978b) |
| CHOP780203 | Normalized frequency of beta-turn | Chou–Fasman (1978b) |
| EISD860101 | Solvation free energy | Eisenberg–McLachlan (1986) |
| FASG760101 | Molecular weight | Fasman (1976) |
| FAUJ830101 | Hydrophobic parameter $\pi$ | Fauchere–Pliska (1983) |
| GRAR740102 | Polarity | Grantham (1974) |
| GRAR740103 | Volume | Grantham (1974) |
| GUYH850101 | Partition energy | Guy (1985) |
| HOPT810101 | Hydrophilicity value | Hopp–Woods (1981) |
| JANJ780101 | Average accessible surface area | Janin et al. (1978) |
| KARP850101 | Flexibility parameter for no rigid neighbors | Karplus–Schulz (1985) |
| KYTJ820101 | Hydropathy index | Kyte–Doolittle (1982) |
| ROSM880101 | Side chain hydropathy (uncorrected for solvation) | Roseman (1988) |
| VINM940101 | Normalized flexibility parameters (B-values), average | Vihinen et al. (1994) |
| WERD780101 | Propensity to be buried inside | Wertz–Scheraga (1978) |
| ZIMJ680101 | Hydrophobicity | Zimmerman et al. (1968) |
| ZIMJ680104 | Isoelectric point | Zimmerman et al. (1968) |

**Table S1: AAindex-derived physicochemical descriptors used for residue encoding.**
This table lists the 20 AAindex properties selected to construct the per-residue physicochemical feature vector. Each property is referenced by its AAindex ID and literature source, and values are z-score normalized across the 20 amino acids prior to model input.

| pHLA | Usage | #HLA | #Peptide | Description | Compared methods |
|---|---|---|---|---|---|
| *Chu et al.* Dataset | Train | 112 | 1,855,306 | Training data | TransPHLA, NetMHCpan-4.1, PickPocket, ANN, DeepAttentionPan, Consensus, ACME |
| | ID test | 112 | 158,145 | In-distribution | |
| | DS test | 5 | 100,592 | distribution-shifted | |
| *Que et al.* Dataset | Train | 95 | 232,163 | Training data | deepAntigen, TransPHLA, NetMHCpan-4.1, MixMHCPred |
| | ID test | 95 | 67,592 | In-distribution | |
| | OOD test | 85 | 26,974 | OOD & zero-shot | |
| | TESLA | 13 | 605 | OOD & zero-shot, tumor neoantigen, TESLA | deepAntigen |
| *Albert et al.* Dataset | Train | 180 | 15,665,005 | Training data | NetMHCpan-4.1, BigMHC, TransPHLA, MixMHCPred |
| | ID test | 36 | 935,497 | In-distribution | |
| | CEDAR | 12 | 1,449 | OOD & zero-shot, tumor neoantigen, CEDAR | NetMHCpan-4.1 |
| Final Dataset | Train | 180 | 16,528,056 | Final training data | / |

**Table S2: Datasets used for training and evaluation of StriMap pHLA predictor.**
Summary of peptide–HLA (pHLA) datasets used to train and evaluate the StriMap pHLA predictor. We refer to in-distribution (ID) evaluation as testing on examples drawn from the same distribution as training; distribution-shifted (DS) evaluation as testing on sets drawn from different studies, cohorts, or experimental conditions; and out-of-distribution (OOD) evaluation as testing on held-out entities such as entirely unseen epitopes and unseen HLAs (peptide and HLA absent from training). Chu et al., Que et al. and Albert et al. datasets were used for benchmarking StriMap against state-of-the-art pHLA prediction methods under ID, DS, and OOD settings, respectively. Final dataset was used for comprehensive training of the final StriMap pHLA models.

| TCR-pHLA | Usage | #HLA | #Peptide | #TCRa | #TCRb | Description | Compared methods |
|---|---|---|---|---|---|---|---|
| *Zhang et al.* Dataset | Train | 19 | 179 | 7,883 | 8,255 | Training data | EPACT, ERGO-AE, NetTCR2.2, STAPLER, MixTCRpred |
| | ID test | 12 | 62 | 1,114 | 1,087 | In-distribution | |
| *Minervina et al.* Dataset | DS test | 5 | 14 | 19,080 | 20,578 | SARS-COV-2 | |
| *Lu et al.* Dataset | Train | 26 | 57 | 4,292 | 4,292 | Training data | TCRconv, TCRGP, TCR-BERT, TCR-H, TEPCAM, EPI-TCR, TEIM, epiTCR, ESM2+LR |
| | ID test | 13 | 18 | 327 | 327 | In-distribution | |
| | OOD test | 56 | 192 | 689 | 689 | Epitope zero-shot | |
| IMMREP22 | Train | 7 | 17 | 6,603 | 6,841 | IMMREP22 challenge | MixTCRpred, TCRAI ERGO2.0, tcrdist3 TCRbase, TCRGP NetTCR2.0, TCRex, |
| | ID test | 7 | 17 | 2,697 | 2,761 | IMMREP22 challenge | |
| IMMREP23 | Train | 39 | 807 | 8,649 | 8,992 | IMMREP23 challenge | IMW DETECT, MixTCRpred, NetTCR, ESM2-shallow, TCRbase |
| | ID test | 6 | 17 | 616 | 607 | IMMREP23 challenge | |
| | OOD test | 1 | 3 | 174 | 176 | IMMREP23 challenge, Epitope zero-shot | |
| Final Dataset | Train | 56 | 1,133 | 17,245 | 17,536 | Final training data | / |
| TP53 Dataset | OOD test | 6 | 6 | 761 | 767 | Epitope zero-shot, Cancer neoantigen | TAPIR, NetTCR2.0 |
| Melanoma Dataset | OOD test | 19 | 6,327 | 94 | 94 | Epitope zero-shot, Cancer neoantigen | / |
| AS Dataset | OOD test | 1 | 13,068,902 | 13 | 7 | Epitope zero-shot, Microbial epitopes for AS | NetTCR-2.2, Logistic regression, Amino Acid Frequency |

**Table S3: Datasets used for training and evaluation of StriMap TCR-pHLA predictor.** Summary of TCR–peptide–HLA (TCR–pHLA) datasets used to train and evaluate the StriMap TCR–pHLA predictor. We refer to in-distribution (ID) evaluation as testing on examples drawn from the same distribution as training under the original split; distribution-shifted (DS) evaluation as testing on sets drawn from different studies, cohorts, or experimental conditions; and out-of-distribution (OOD) evaluation as testing on held-out entities such as entirely unseen epitopes (peptide sequences absent from training). *Zhang et al.*, *Minervina et al.*, *Lu et al.*, IMMREP22 and IMMREP23 datasets were used for benchmarking StriMap against existing TCR–pHLA prediction methods across multiple scenarios, including ID, DS, and OOD. Final dataset was used for comprehensive training of the final StriMap TCR–pHLA models. TP53 and Melanoma Datasets were used for cancer neoantigen analyses, and AS Dataset was used for ankylosing spondylitis–associated microbial epitope analysis.

| Species | Strain ID | Reference | #Proteins |
|---|---|---|---|
| *Klebsiella pneumoniae* | RJX1596 | PMID: 29438717,36477533 | 5562 |
| *Escherichia coli* | RJX1588 | PMID: 37494472,33986841,36477533 | 4692 |
| *Ruminococcus gnavus* | RJX1119 | PMID: 37494472,35818337 | 2994 |
| *Streptococcus anginosus (1)* | RJX1754 | PMID: 37494472,35794662 | 442 |
| *Streptococcus anginosus (2)* | J4211 | PMID: 37494472,35794662 | 1926 |
| *Streptococcus mutans* | NCH105 | PMID: 37494472,35794662 | 1911 |
| *Streptococcus parasanguinis* | RJX1181 | PMID: 37494472,35794662 | 1953 |
| *Streptococcus salivarius* | Ssal_L25 | PMID: 37494472,35794662 | 1919 |
| *Akkermansia muciniphila* | JCM 30893 | PMID: 37494472,25434931 | 2375 |
| *Clostridium bartlettii / Intestinibacter bartlettii* | MGYG-HGUT-00062 | PMID: 37494472 | 2860 |
| *Lactobacillus fermentum / Limosilactobacillus fermentum* | EFEL6800 | | 1991 |
| *Lachnospiraceae bacterium* | UHGG_MGYG-HGUT-02492 | | 3077 |
| *Parasutterella excrementihominis (1)* | RJX1996 | | 3639 |
| *Parasutterella excrementihominis (2)* | YIT 11859 | | 2751 |
| *Prevotella copri / Segatella copri* | DSM 18205 | PMID: 28750650 | 2853 |
| *Prevotella melaninogenica* | ATCC 25845 | PMID: 28750650 | 2296 |

**Table S4: Gut bacterial species associated with ankylosing spondylitis used for large-scale peptide screening.**
List of bacterial strains reported to be enriched or associated with ankylosing spondylitis (AS) or related inflammatory conditions based on prior metagenomic and clinical studies. For each strain, corresponding literature references (PMIDs) and the total number of annotated protein-coding sequences used for peptide enumeration are shown. All possible 9-mer peptides derived from these proteins were included in downstream HLA-B*27:05 binding prediction and TCR–pHLA screening analyses.

| kruskalTestP | comp | Z | padj | var | Layer |
|---|---|---|---|---|---|
| 4.97E-29 | Healthy - CD | -8.9700867 | 2.96E-18 | k119_53774_1_3 | MGX |
| 4.97E-29 | Healthy - UC | -3.2556341 | 0.01131395 | k119_53774_1_3 | MGX |
| 3.98E-83 | Healthy - CD | -3.9627107 | 0.00074104 | k141_49810_3 | MGX |
| 3.98E-83 | Healthy - Gastroenteritis | -17.852198 | 2.78E-70 | k141_49810_3 | MGX |
| 0.01675648 | Healthy - UC | 2.83751466 | 0.04546626 | k97_1417670_1 | MGX |
| 2.31E-41 | Healthy - CD | -8.7994335 | 1.38E-17 | k97_1449468_1 | MGX |
| 2.31E-41 | Healthy - Gastroenteritis | 4.10790962 | 0.00039926 | k97_1449468_1 | MGX |
| 2.31E-41 | Healthy - UC | -4.4823247 | 7.38E-05 | k97_1449468_1 | MGX |
| 1.97E-07 | Healthy - CD | -5.1011553 | 3.38E-06 | k97_1999973_2 | MGX |
| 0.00945233 | Healthy - Gastroenteritis | 2.9469594 | 0.03209153 | k97_4911109_16 | MGX |
| 0.00945233 | Healthy - UC | 2.52253191 | 0.11651338 | k97_4911109_16 | MGX |
| 2.30E-56 | Healthy - CD | -5.8458836 | 5.04E-08 | k99_11015_5_2 | MGX |
| 2.30E-56 | Healthy - Gastroenteritis | -15.057841 | 3.07E-50 | k99_11015_5_2 | MGX |
| 5.57E-16 | Healthy - CD | -4.6646682 | 3.09E-05 | k99_24291_10 | MGX |
| 5.57E-16 | Healthy - UC | -6.3089054 | 2.81E-09 | k99_24291_10 | MGX |
| 7.56E-10 | Healthy - UC | -6.2648845 | 3.73E-09 | k99_24291_11 | MGX |
| 1.02E-14 | Healthy - CD | -4.6144115 | 3.94E-05 | k99_24291_15 | MGX |
| 1.02E-14 | Healthy - UC | -6.2544035 | 3.99E-09 | k99_24291_15 | MGX |
| 4.16E-10 | Healthy - CD | -3.169185 | 0.01528671 | k99_24291_18 | MGX |
| 4.16E-10 | Healthy - UC | -5.2280146 | 1.71E-06 | k99_24291_18 | MGX |
| 4.10E-25 | Healthy - Gastroenteritis | 7.52504495 | 5.27E-13 | k99_71711_2_2 | MGX |
| 4.10E-25 | Healthy - UC | 3.05438695 | 0.0225521 | k99_71711_2_2 | MGX |
| 0.04200476 | Healthy - CD | -3.0096624 | 0.02615382 | k99_71711_2_3 | MGX |
| 2.39E-11 | Healthy - CD | -6.4621982 | 1.03E-09 | k99_81598_2_4 | MGX |
| 8.70E-31 | Healthy - Gastroenteritis | 11.120438 | 9.98E-28 | k99_71711_2 | MTX |
| 8.70E-31 | Healthy - UC | 2.28724182 | 0.22181713 | k99_71711_2 | MTX |
| 1.62E-16 | Healthy - Gastroenteritis | 8.29986495 | 1.04E-15 | k97_4911109_16 | MTX |
| 1.62E-16 | Healthy - UC | 3.45818341 | 0.00543831 | k97_4911109_16 | MTX |
| 4.65E-05 | Healthy - CD | -3.6194709 | 0.00295206 | k97_1999973_2 | MTX |
| 4.65E-05 | Healthy - UC | -3.3398457 | 0.00838249 | k97_1999973_2 | MTX |
| 3.53E-17 | Healthy - Gastroenteritis | 7.05149213 | 1.77E-11 | k99_34556_2 | MTX |
| 1.06E-10 | Healthy - CD | -5.7792618 | 7.50E-08 | k99_81598_2_4 | MTX |
| 5.07E-06 | Healthy - CD | -2.5482831 | 0.10825459 | k99_71711_2_2 | MTX |
| 5.07E-06 | Healthy - UC | -2.5202803 | 0.11726143 | k99_71711_2_2 | MTX |
| 1.21E-06 | Healthy - CD | -3.880142 | 0.00104395 | k99_75096_12 | MTX |
| 1.21E-06 | Healthy - UC | -3.1068675 | 0.01890811 | k99_75096_12 | MTX |
| 1.71E-104 | Healthy - CD | -14.958808 | 1.36E-49 | k97_1449468_1 | MTX |
| 1.71E-104 | Healthy - UC | -11.381755 | 5.16E-29 | k97_1449468_1 | MTX |
| 1.24E-20 | Healthy - CD | 2.30415065 | 0.21214188 | k119_19123_9 | MTX |
| 1.24E-20 | Healthy - Gastroenteritis | 7.37140585 | 1.69E-12 | k119_19123_9 | MTX |

**Table S5: Differential abundance/expression of candidate peptide–encoding features across cohorts (MGX/MTX).**

This table summarizes nonparametric group comparisons for each feature (var) across disease cohorts and assay layers. kruskalTestP reports the overall Kruskal-Wallis test p value for a given feature across all cohorts. comp indicates the specific pairwise comparison (e.g., Healthy – CD,

Healthy – UC, Healthy – Gastroenteritis). Z is the standardized test statistic from post hoc pairwise testing (e.g., Dunn's test), where the sign reflects the direction of change relative to Healthy (negative Z indicates higher values in the non-Healthy group when the comparison is defined as Healthy minus group). padj is the multiple-testing–adjusted p value for the pairwise comparison. var denotes the feature identifier (e.g., protein ID). Layer indicates the data modality: MGX (shotgun metagenomics) and MTX (metatranscriptomics).

### Supplementary Figures

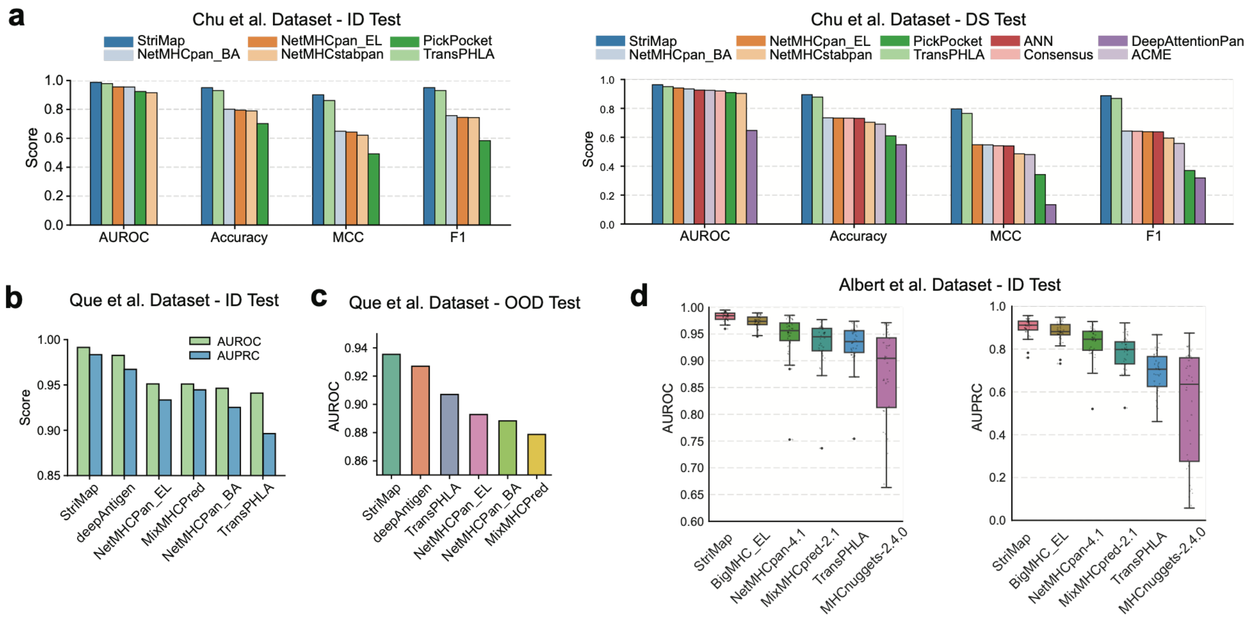


**Fig. S1: Extended benchmarking results for pHLA binding prediction.**
**(a)** Performance on the *Chu et al.* dataset under ID and DS test settings measured by AUROC, Accuracy, MCC, and F1 score.
**(b)** AUROC and AUPRC comparison on the *Que et al.* dataset under the ID setting.
**(c)** AUROC comparison on the *Que et al.* dataset under the OOD setting.
**(d)** Boxplots of AUROC and AUPRC on the *Albert et al.* dataset under the ID setting. Center lines indicate medians, boxes represent interquartile ranges, and whiskers denote 1.5× the interquartile range.

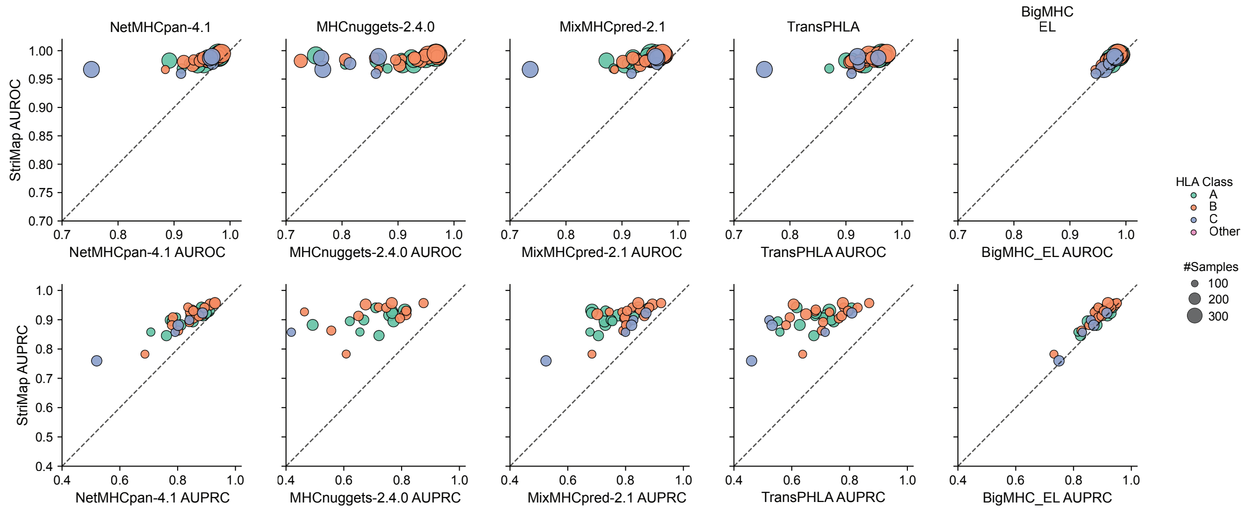


**Fig. S2: Per-allele performance comparison of StriMap and state-of-the-art pHLA binding predictors.**

Per-HLA AUROC (top row) and AUPRC (bottom row) comparisons between StriMap and five leading pHLA prediction models: NetMHCpan-4.1, MHCnuggets-2.4.0, MixMHCpred-2.1, TransPHLA, and BigMHC-EL. Each point represents a single HLA allele. Colors denote HLA class (A, B, C, or other), and point sizes indicate the number of peptides associated with each allele. The dashed diagonal lines indicate equal performance between StriMap and the corresponding comparison method.

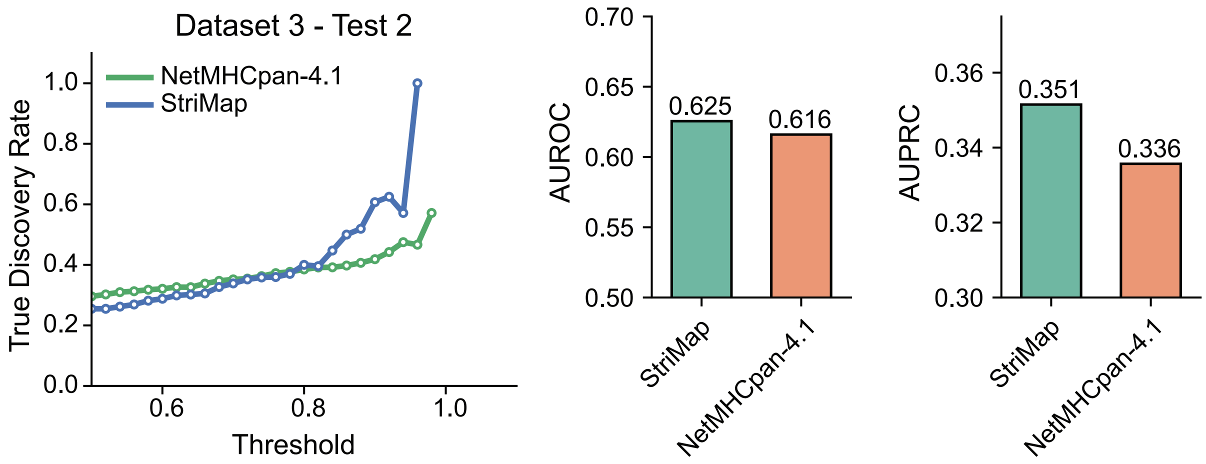


**Fig. S3: Performance comparison of StriMap and NetMHCpan-4.1 on the CEDAR benchmark.**

StriMap and NetMHCpan-4.1 were evaluated on the CEDAR dataset (Dataset 3 Test 2) using the same train–test split defined in the original NetMHCpan study. Left, true discovery rate (TDR) as a function of ranking threshold. Middle, overall AUROC. Right, overall AUPRC.

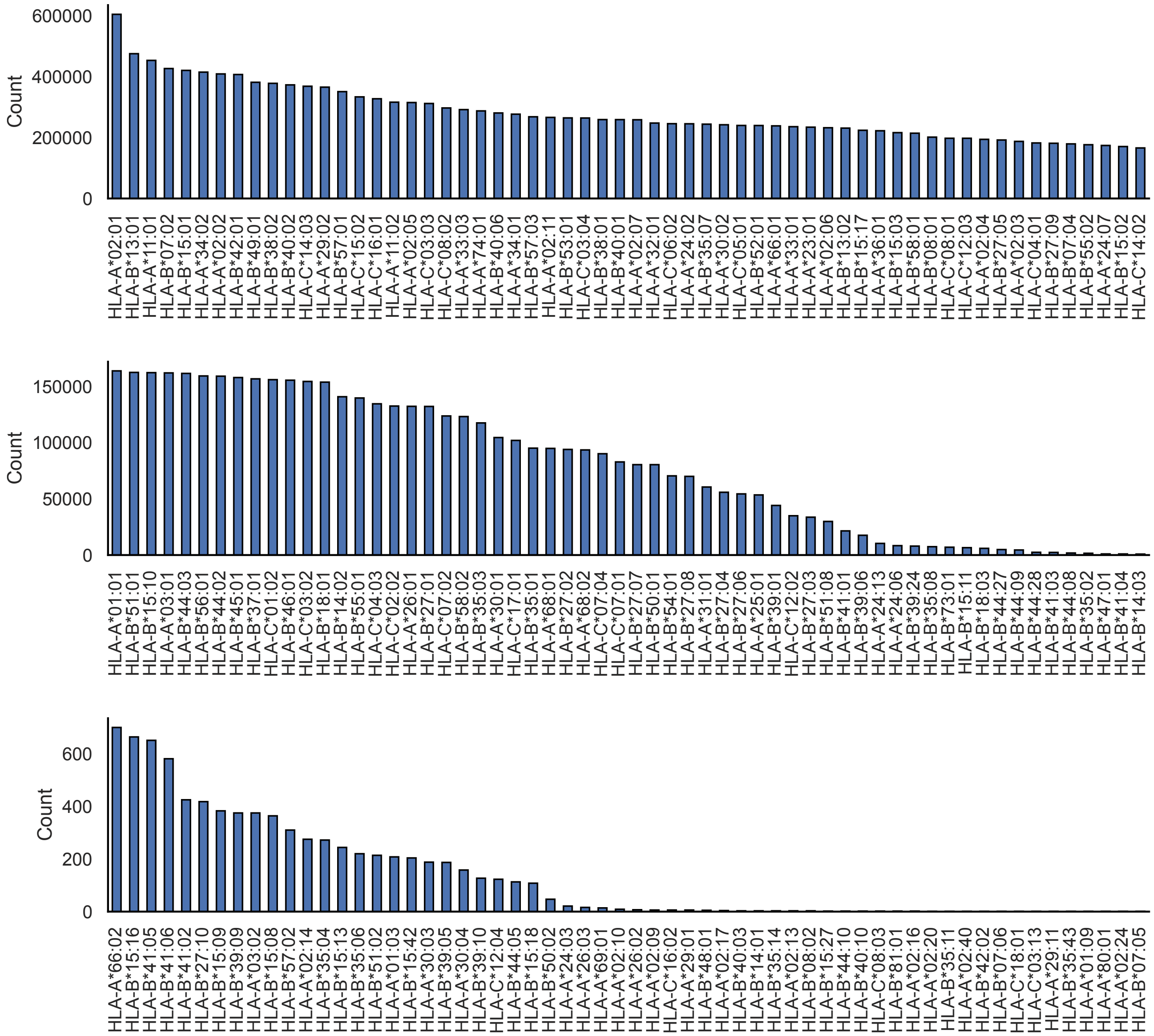


**Fig. S4: Distribution of peptide counts across HLA alleles in the final training dataset.** Bar plots show the number of peptides associated with each HLA allele in the final training dataset, stratified by HLA class. HLA alleles are ordered by decreasing peptide count within each panel, illustrating the coverage and imbalance of HLA representation in the training data.

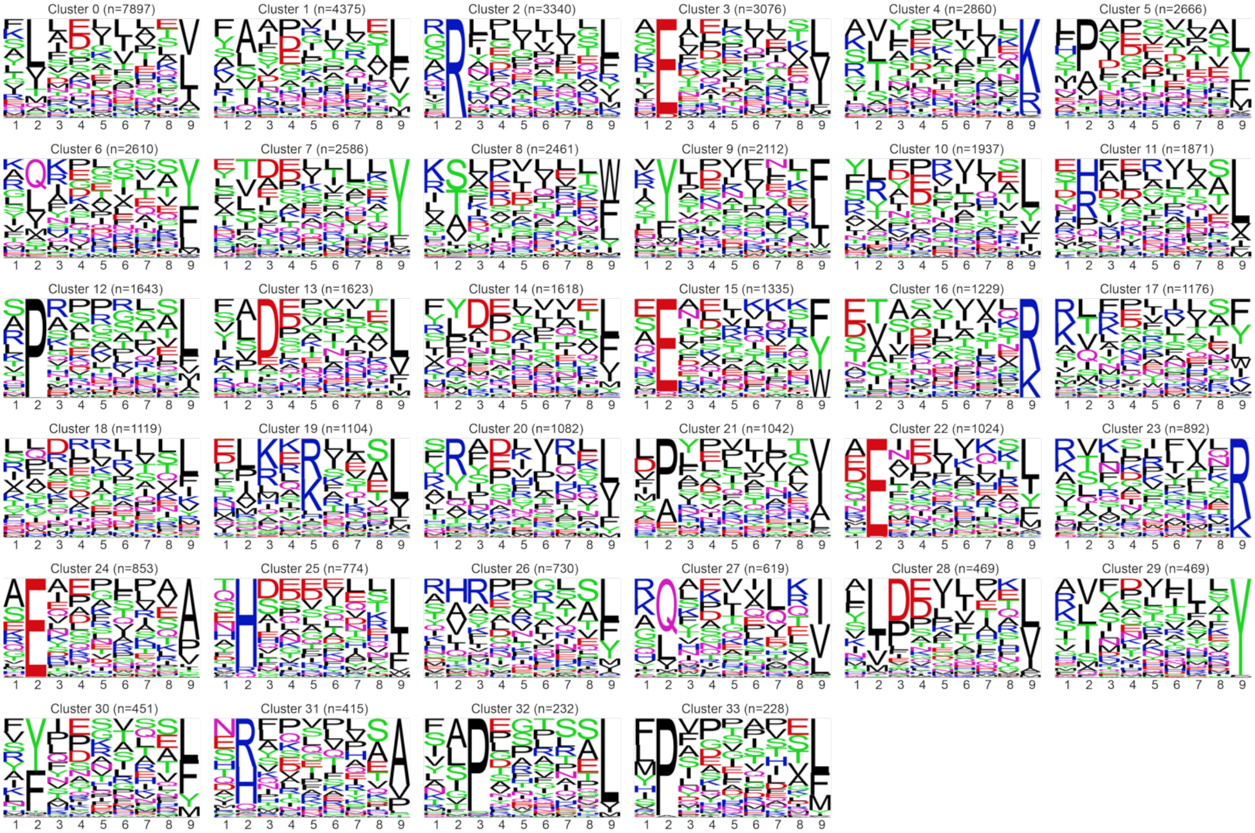

**Fig. S5: Peptide sequence motifs of Leiden clusters identified in Fig. 2d.**
Sequence logos representing peptide sequence motifs for each Leiden cluster shown in Fig. 2d. For each cluster, logos summarize position-specific amino acid preferences of 9-mer peptides, highlighting distinct binding motifs associated with different peptide–HLA interaction patterns.

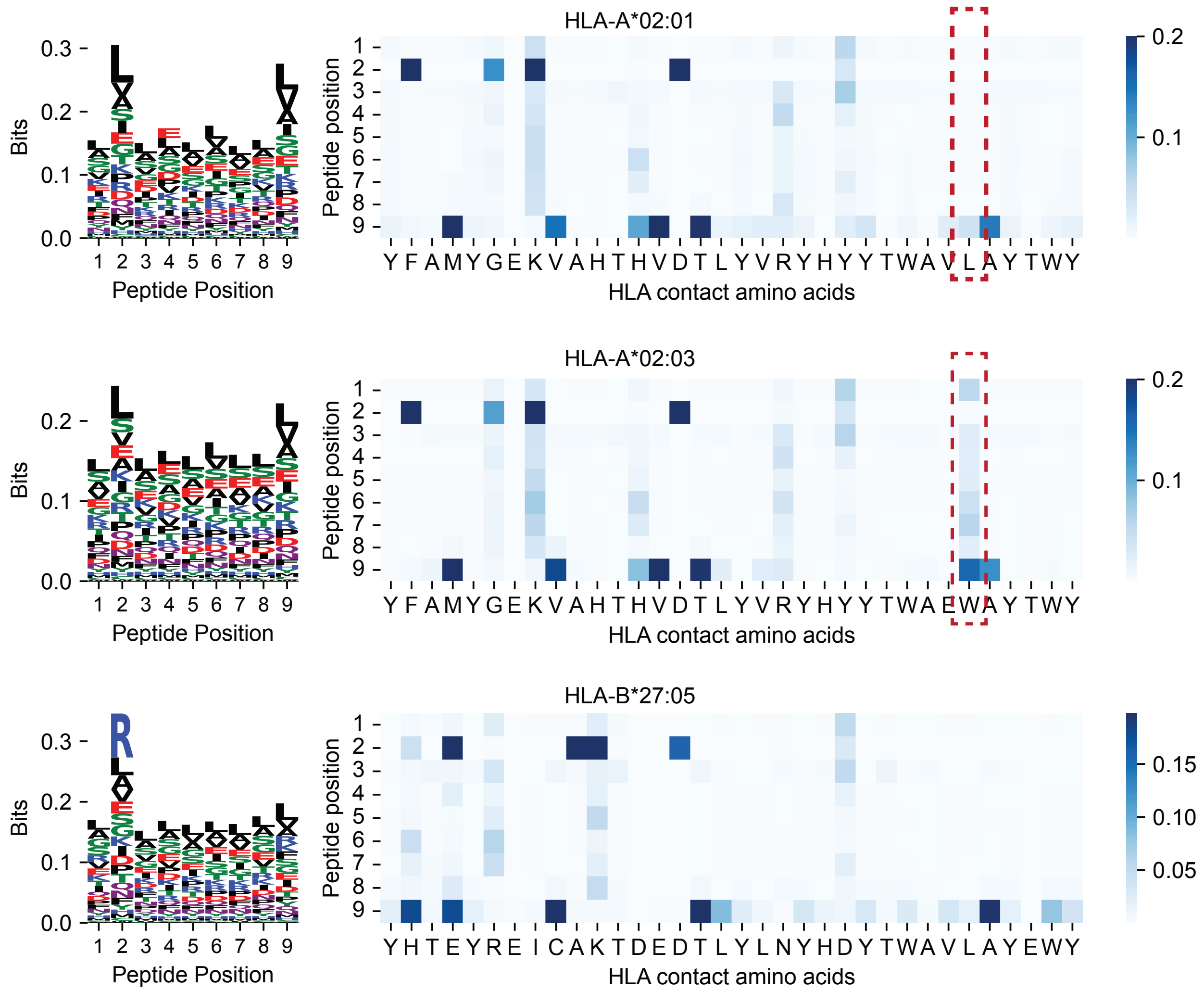


**Fig. S6: Attention patterns underlying peptide–HLA binding learned by StriMap.** Sequence logos (left) depict position-specific amino acid preferences of 9-mer peptides for representative HLA alleles, illustrating canonical peptide-binding motifs. Heatmaps (right) show normalized attention weights between peptide positions (rows) and HLA contact amino acids (columns) learned by the StriMap model. Results are shown for HLA-A02:01, HLA-A02:03, and HLA-B*27:05, highlighting allele-specific interaction patterns between peptide residues and HLA binding pockets.

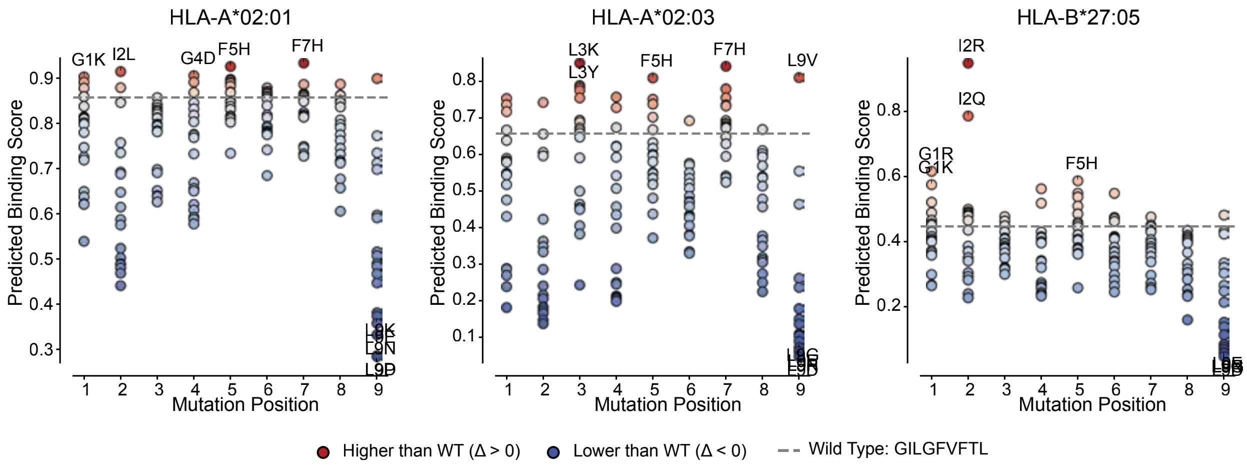


**Fig. S7: Saturation mutagenesis analysis of the influenza peptide GILGFVFTL across representative HLA alleles.**

Saturation mutagenesis analysis of the influenza A–derived peptide GILGFVFTL, showing predicted binding scores for all single–amino acid substitutions across peptide positions for HLA-A*02:01, HLA-A*02:03, and HLA-B*27:05. Each point represents a single mutant peptide. Colors indicate substitutions predicted to increase (red) or decrease (blue) binding affinity relative to the wild-type peptide (gray dashed line). Selected mutations with pronounced effects are annotated, illustrating position- and allele-specific sensitivity of peptide–HLA binding learned by the StriMap model.

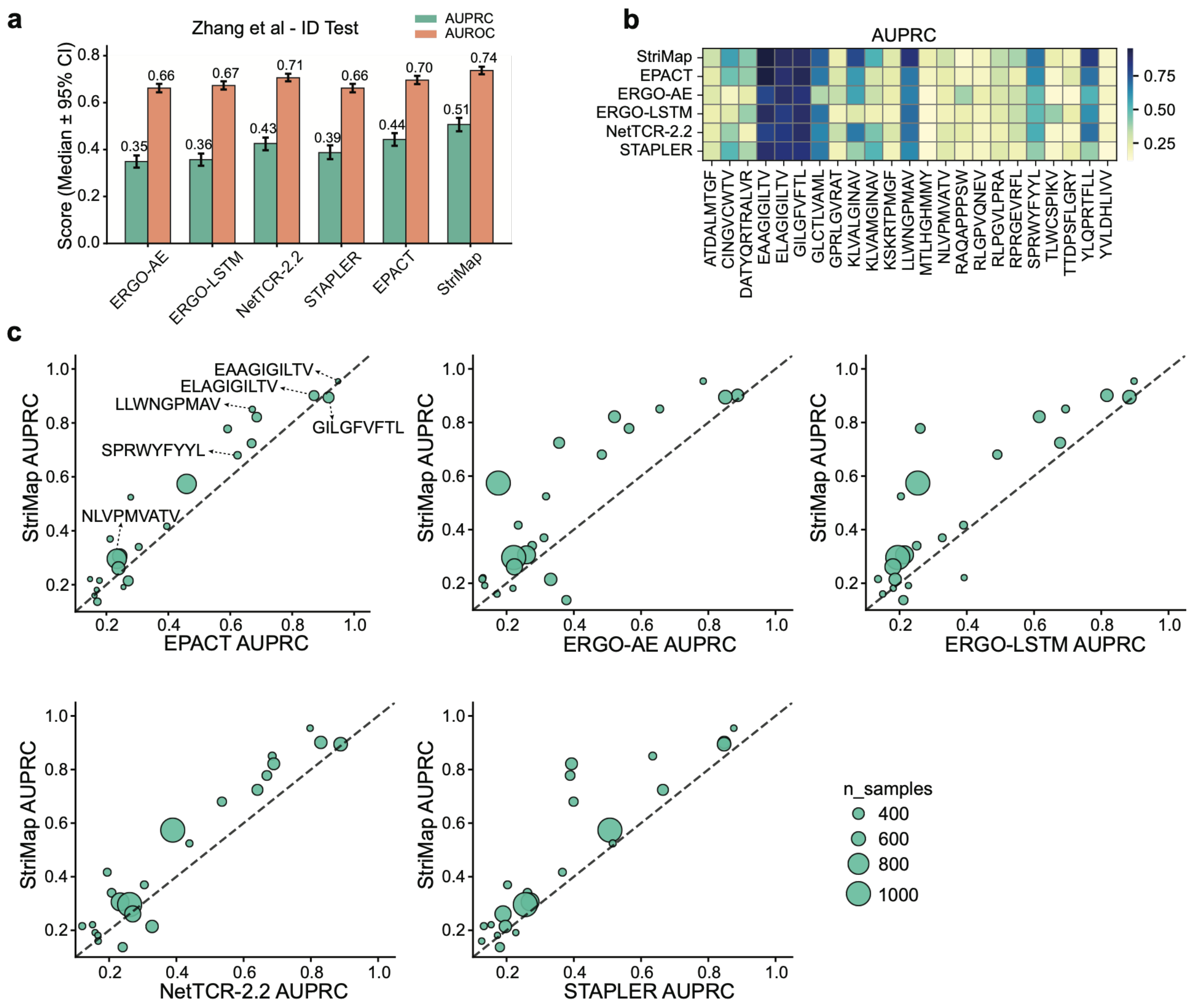


**Fig. S8: Per-peptide performance analysis on the Zhang et al. dataset under the in-distribution (ID) test setting.**

**(a)** Overall performance comparison on the Zhang et al. dataset under the ID test setting. Models are evaluated using AUPRC and AUROC. Bars indicate median performance across runs, with error bars denoting 95% confidence intervals.

**(b)** Heatmap of AUPRC values for individual peptide–HLA pairs across different prediction methods. Rows correspond to models and columns correspond to peptides. Color intensity reflects AUPRC, highlighting variability in per-peptide performance across methods.

**(c)** Pairwise comparison of per-peptide AUPRC between StriMap and baseline methods (EPACT, ERGO-AE, ERGO-LSTM, NetTCR-2.2, and STAPLER). Each point represents a peptide–HLA pair, with point size proportional to the number of samples associated with that pair. The dashed diagonal indicates equal performance between StriMap and the corresponding baseline

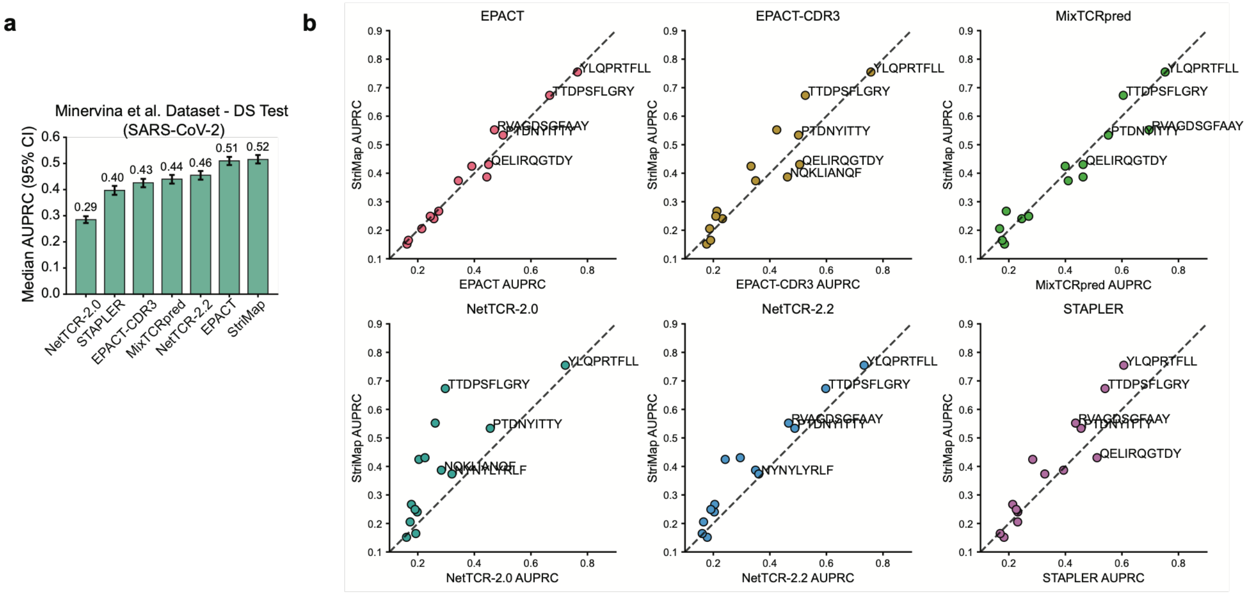


**Fig. S9: Performance comparison on the Minervina et al. dataset under distribution-shift (DS) conditions.**

**(a)** Overall performance comparison on the Minervina et al. dataset under the distribution-shift (DS) test setting, where SARS-CoV-2–related TCR–pMHC interactions are evaluated. Model performance is measured using AUPRC. Bars indicate median AUPRC values across runs, with error bars denoting 95% confidence intervals.

**(b)** Pairwise comparison of per-peptide AUPRC between StriMap and baseline methods, including EPACT, EPACT-CDR3, MixTCRpred, NetTCR-2.0, NetTCR-2.2, and STAPLER. Each point represents a peptide–HLA pair. The dashed diagonal indicates equal performance between StriMap and the corresponding baseline, with points above the diagonal indicating improved performance by StriMap.

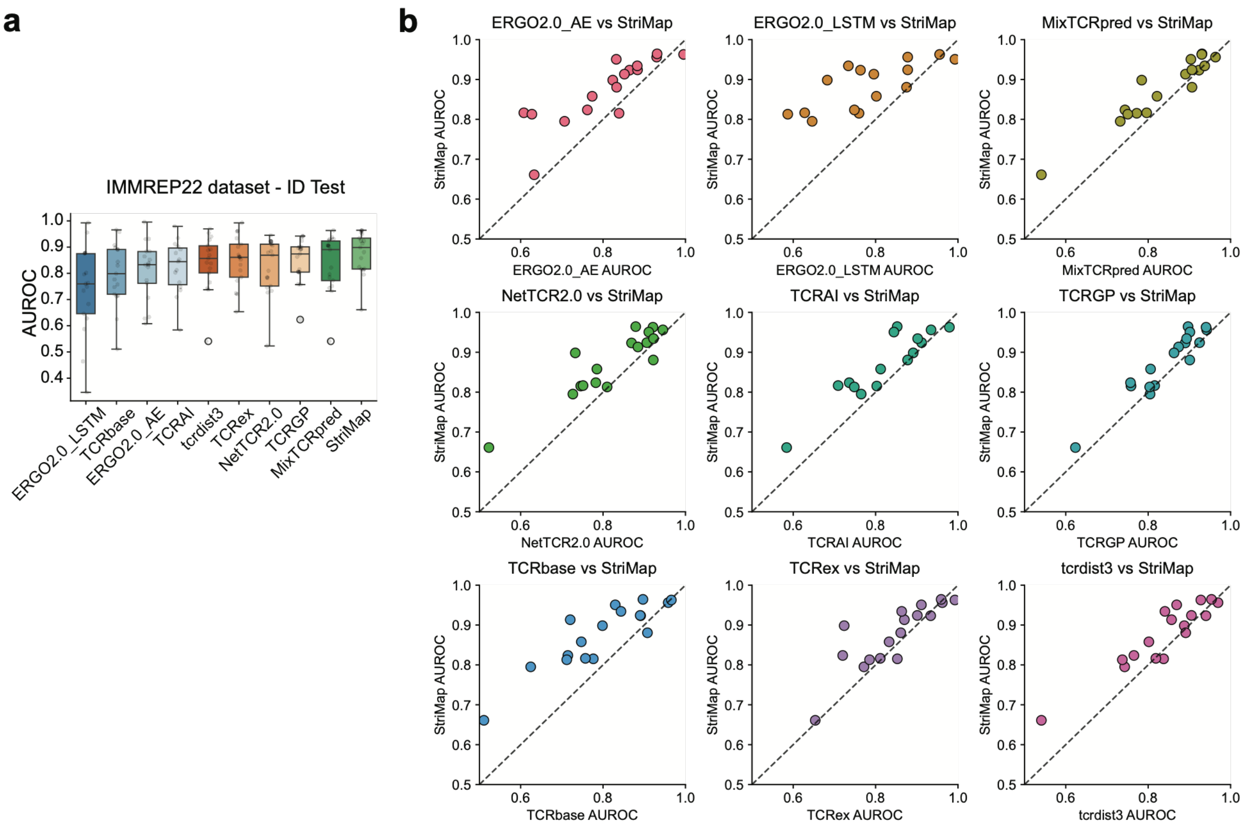


**Fig. S10: Performance comparison on the IMMREP22 dataset under the in-distribution (ID) test setting.**

**(a)** Overall performance comparison on the IMMREP22 dataset under the ID test setting, evaluated using AUROC. Boxplots summarize the distribution of AUROC values across multiple runs or data splits for each method. Center lines indicate medians, boxes represent interquartile ranges, and whiskers denote 1.5× the interquartile range.

**(b)** Pairwise comparison of AUROC between StriMap and baseline methods, including ERGO2.0-AE, ERGO2.0-LSTM, MixTCRpred, NetTCR2.0, TCRAI, TCRGP, TCRbase, TCRex, and tcrdist3. Each point represents a peptide–HLA pair. The dashed diagonal indicates equal performance between StriMap and the corresponding baseline, with points above the diagonal indicating higher AUROC achieved by StriMap.

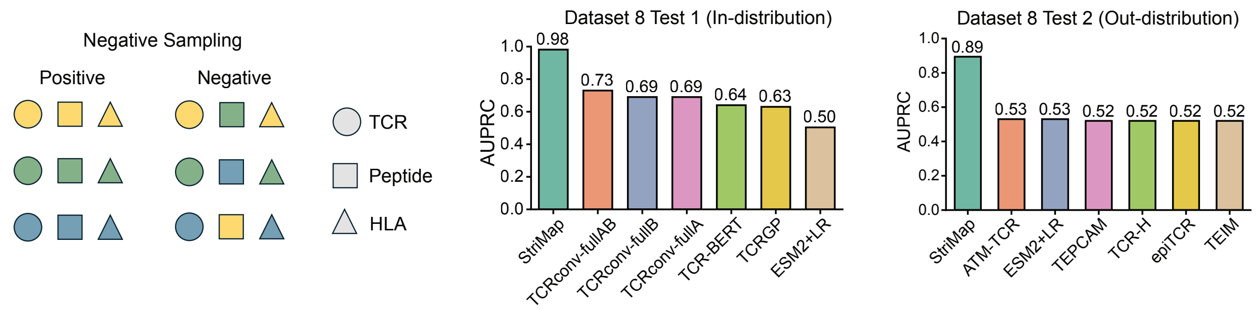


**Fig. S11: Impact of negative sampling strategies on TCR–pHLA prediction benchmarks.** Schematic illustration of the negative sampling strategy used in a recently published benchmark (Lu et al. dataset), in which TCR–HLA pairs are preserved while peptide sequences are randomly permuted to generate negative examples (left). Performance comparison of StriMap and existing TCR–pHLA prediction methods under this evaluation protocol on Dataset 8 Test 1 (independent) and Dataset 8 Test 2 (unseen), measured by AUPRC (right).

**Note: Interpretation of benchmark results**
Under this negative sampling scheme, a large fraction of negative examples can be distinguished based primarily on peptide–HLA compatibility, without requiring accurate modeling of TCR recognition. As StriMap explicitly models peptide–HLA presentation in addition to TCR–pHLA interactions, it achieves substantially higher performance in this setting. In contrast, methods that do not incorporate HLA-dependent peptide presentation are disadvantaged by construction. Consequently, performance differences observed under this evaluation protocol primarily reflect sensitivity to peptide–HLA compatibility rather than TCR-specific recognition accuracy.

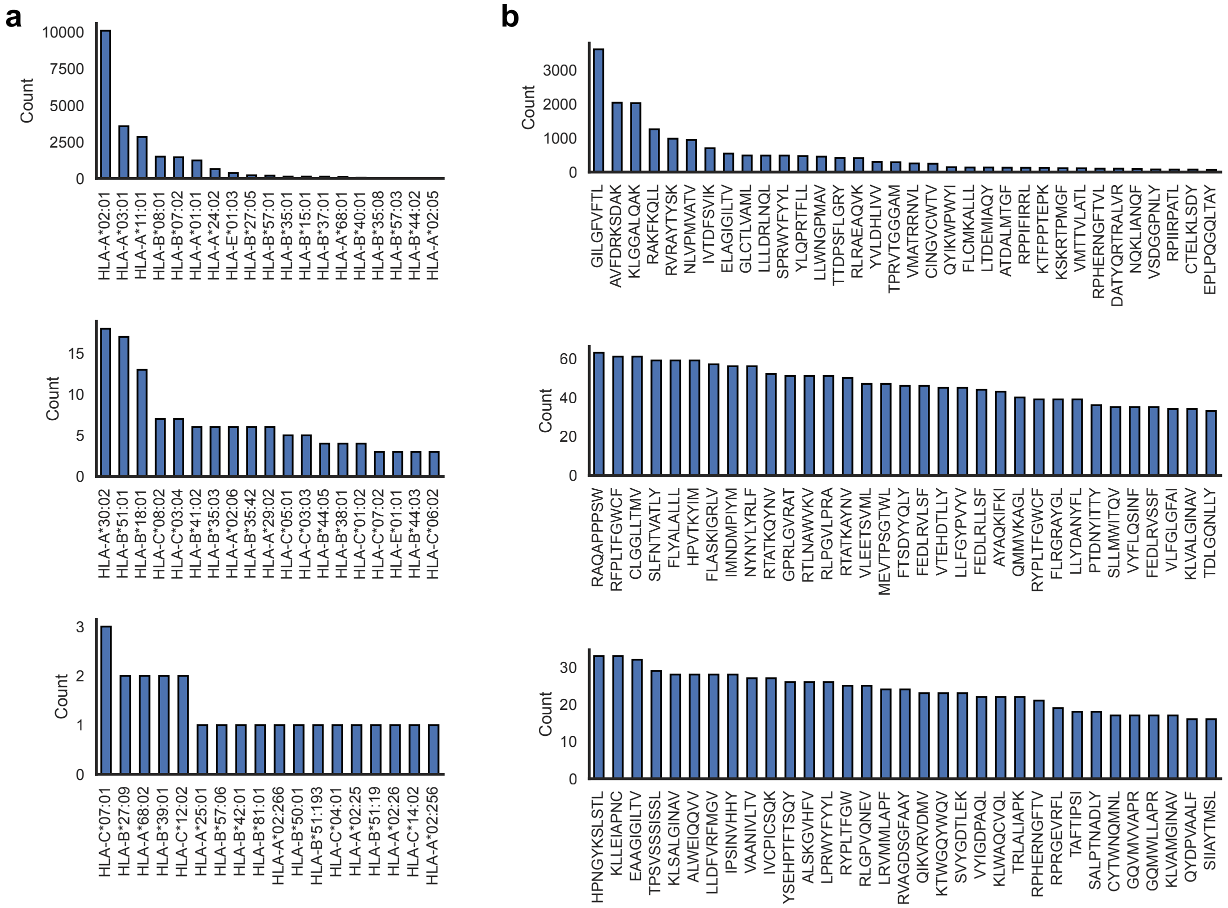


**Fig. S12: Distribution of training data used for the StriMap TCR–pHLA predictor.**
**(a)** Distribution of HLA alleles represented in the final training dataset for the StriMap TCR–pHLA predictor, shown as the number of TCR–peptide pairs associated with each HLA allele.
**(b)** Distribution of peptide sequences in the final training dataset, shown as the number of associated TCRs per peptide. Peptides are ordered by decreasing frequency, illustrating the heterogeneous coverage of peptide antigens in the training data.

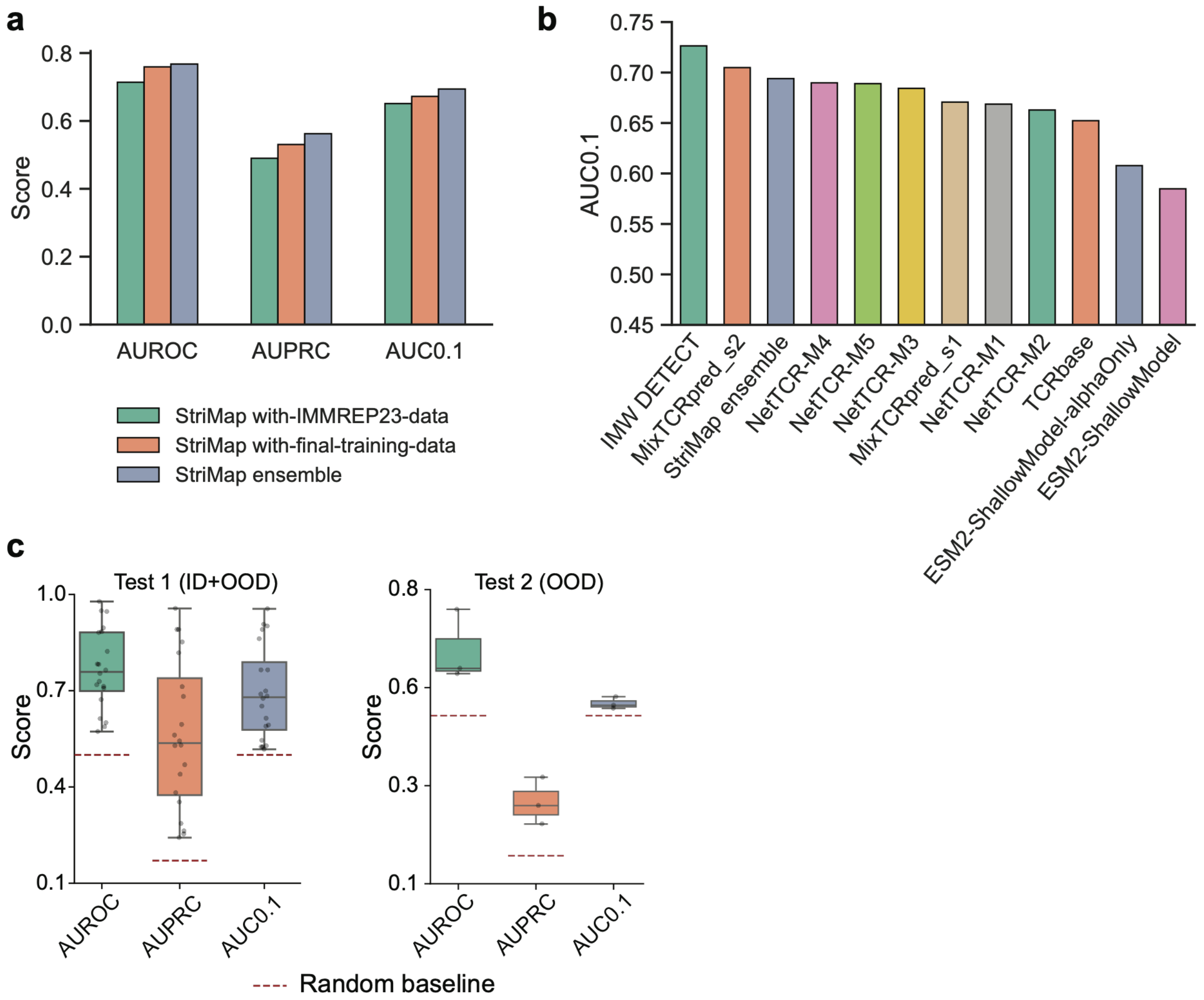


**Fig. S13: Performance of StriMap on the IMMREP23 challenge benchmark.**
**(a)** Performance of StriMap on the IMMREP23 benchmark under different training strategies, evaluated using AUROC, AUPRC, and AUC0.1. Results are shown for models fine-tuned using IMMREP23 data, models fine-tuned using the final training dataset, and an ensemble of StriMap models trained with different negative sampling strategies.
**(b)** AUC0.1 comparison between StriMap and baseline methods evaluated on the IMMREP23 benchmark. StriMap ensemble denotes the aggregation of predictions from multiple StriMap models trained using distinct negative sampling strategies.
**(c)** Distribution of StriMap performance on the IMMREP23 benchmark across evaluation splits for Test 1 (ID+OOD) and Test 2 (OOD), summarized by boxplots for AUROC, AUPRC, and AUC0.1. Center lines indicate medians, boxes represent interquartile ranges, and whiskers denote 1.5× the interquartile range; points correspond to individual splits. Dashed horizontal lines denote the random baseline expected under chance performance for each metric.

Note: IMW DETECT and MixTCRpred_s2 both leveraged undisclosed in-house training data, precluding direct attribution of their performance gains to model architecture versus data availability.

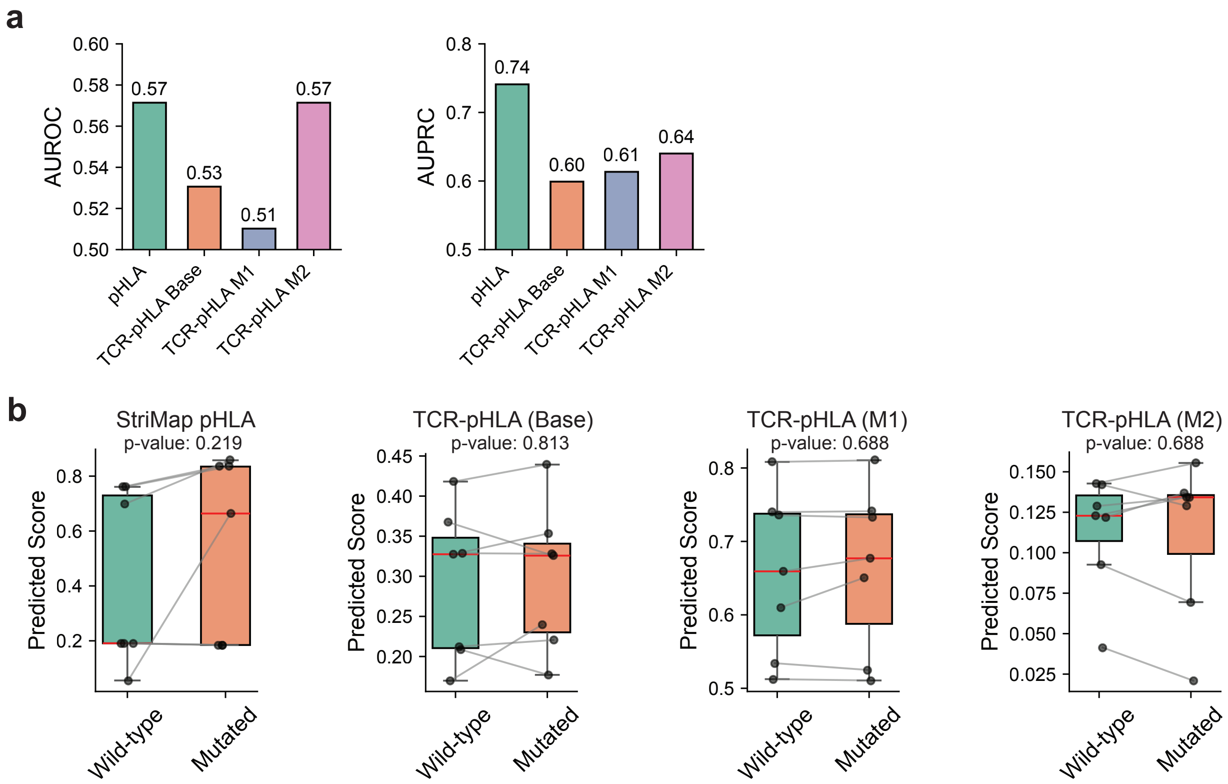


**Fig. S14: StriMap pHLA scoring and TCR–pHLA scoring for wild-type and mutant peptides.**

**(a)** AUROC and AUPRC for distinguishing wild-type from mutant peptides using StriMap pHLA scores or TCR–pHLA scores (wild-type peptides labeled as 1 and mutant peptides labeled as 0). **(b)** Each panel shows paired pHLA scores or TCR-pHLA scores for the same pHLA pair before and after mutation; p values were computed using a paired Wilcoxon signed-rank test.

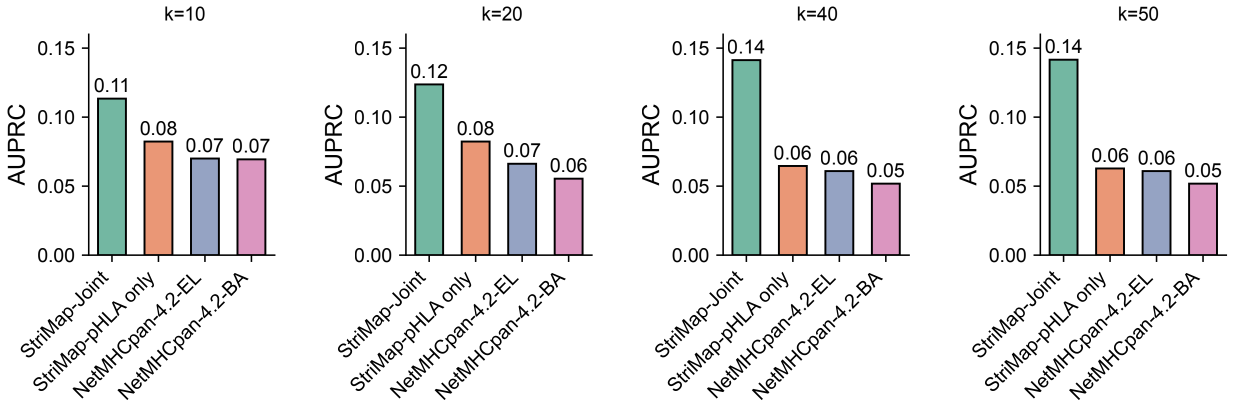


**Fig. S15: Robustness of joint TCR–pHLA ranking across different top-k thresholds.** AUPRC performance of StriMap and baseline methods under varying top-k settings for joint neoepitope prioritization, corresponding to the analysis shown in Fig. 3. Results are shown for k = 10, 20, 40, and 50, where the joint score aggregates the top-k ranked TCR–pHLA interactions for each candidate neoepitope. StriMap-joint consistently outperforms pHLA-only ranking and NetMHCpan-4.2-based baselines across all k values.

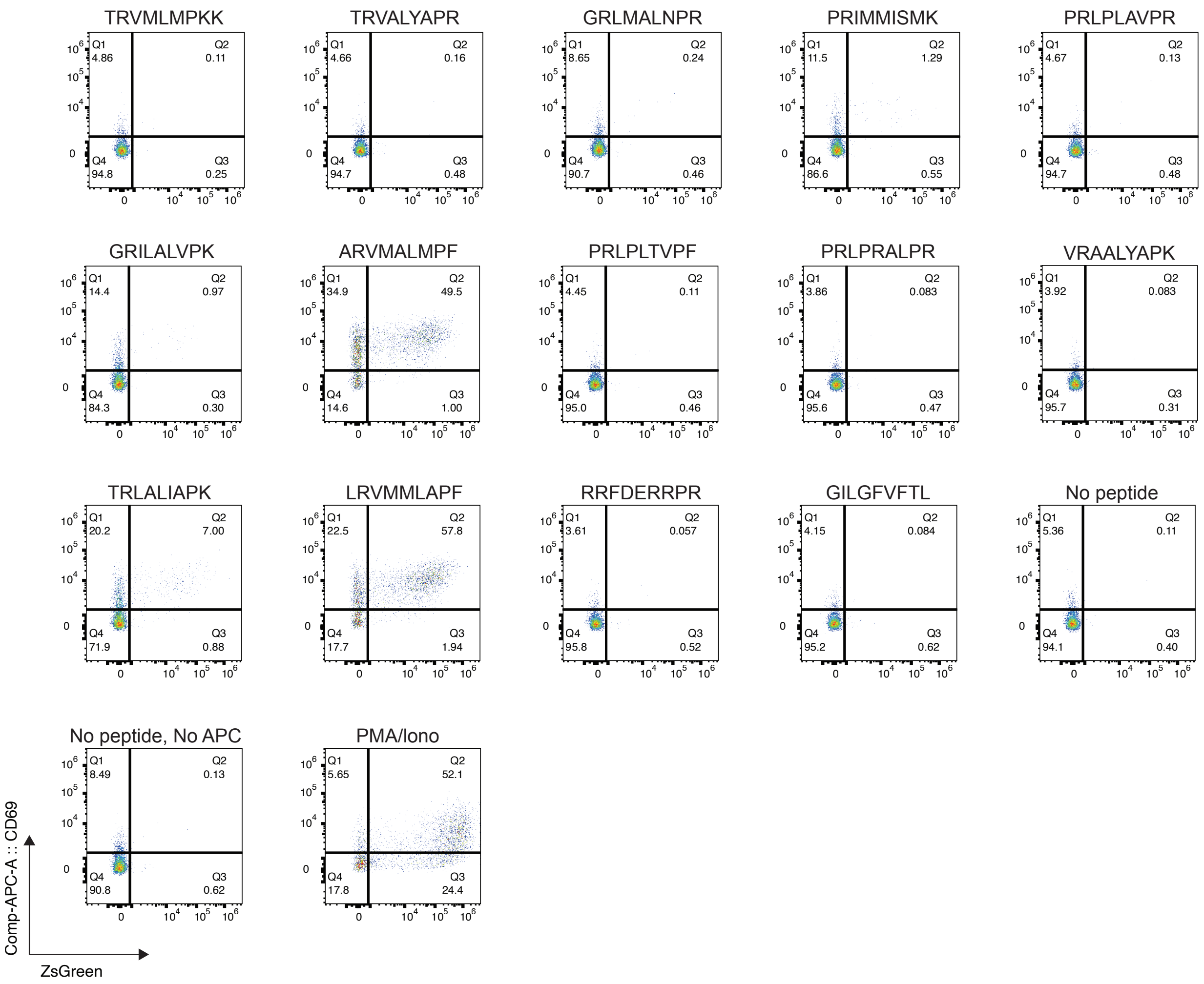


**Fig. S16: Flow cytometry analysis of T cell activation across all tested peptides and control conditions in Fig. 5.**

Representative flow cytometry plots showing CD69 expression (y-axis) and NFAT-driven ZsGreen reporter activity (x-axis) for AS8.4 TCR–expressing Jurkat T cells co-cultured with HLA-B*27:05–expressing antigen-presenting cells pulsed with individual peptide candidates. Each panel corresponds to a single peptide or control condition tested in Fig. 5, including positive hits, non-activating peptides, wild-type peptide controls, and negative controls ("No peptide" and "No peptide, No APC"), as well as PMA/ionomycin stimulation as a positive activation control. Percentages indicate the fraction of cells in each quadrant.

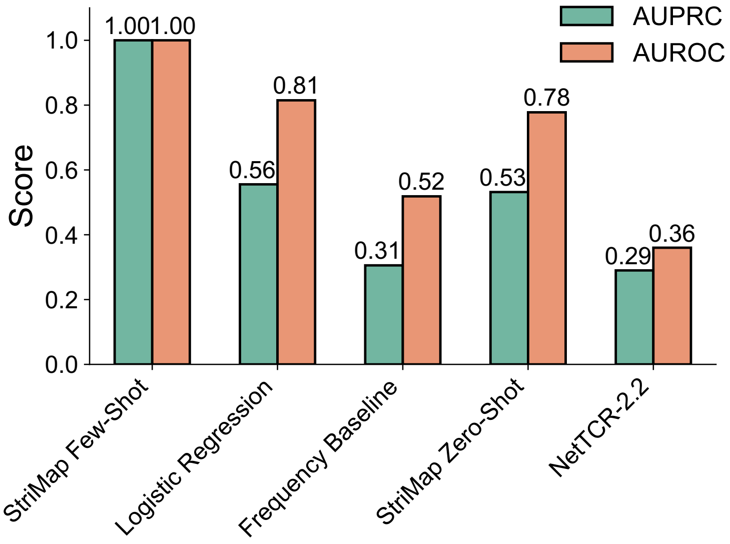


**Fig. S17: Predictive performance of different models for experimentally validated TCR–peptide interactions in Fig. 5.**

Comparison of predictive performance across different modeling strategies for the experimentally tested peptide candidates shown in Fig. 5, evaluated using AUROC and AUPRC. All models, except StriMap in the zero-shot setting and NetTCR-2.2, were trained under the same few-shot regime using limited experimentally validated samples. These include StriMap (few-shot), a logistic regression baseline, and a frequency-based baseline. StriMap-few-shot achieves the highest performance, indicating that incorporating limited experimental feedback substantially improves prediction accuracy for TCR-mediated peptide recognition.

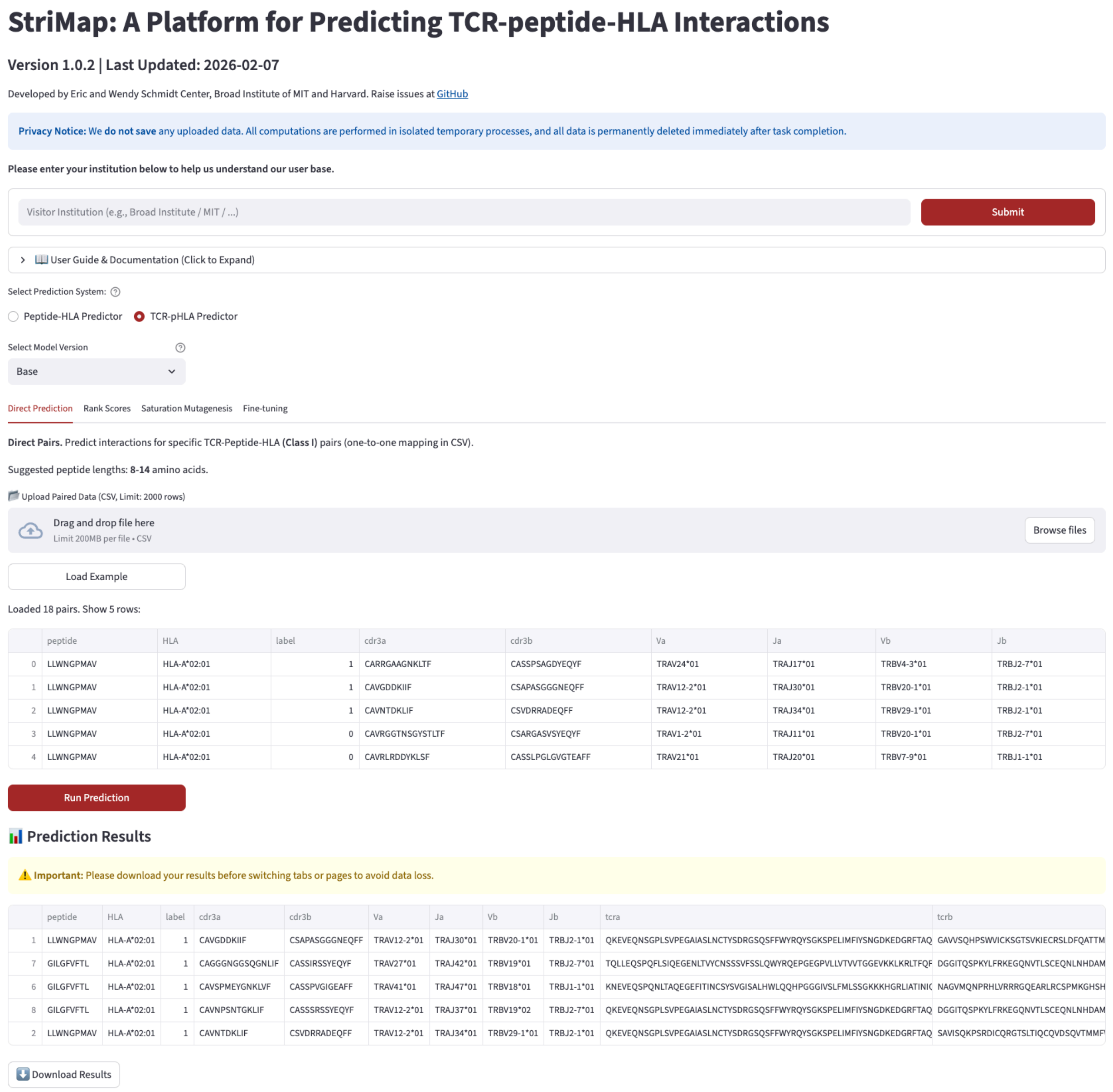


**Fig. S18: Web interface for the StriMap TCR–peptide–HLA predictor.**
Screenshot of the StriMap web platform illustrating the TCR–peptide–HLA prediction interface. Users can upload paired TCR, peptide, and HLA data to perform direct interaction prediction, ranking of candidate interactions, and optional on-the-fly fine-tuning. The platform provides three model variants (Base, M1, and M2), corresponding to different negative sampling strategies used during training. All computations are executed in isolated temporary processes without data retention, enabling secure analysis of sensitive immune receptor data.
